# A COVID-19 vaccine candidate using SpyCatcher multimerization of the SARS-CoV-2 spike protein receptor-binding domain induces potent neutralising antibody responses

**DOI:** 10.1101/2020.08.31.275701

**Authors:** Tiong Kit Tan, Pramila Rijal, Rolle Rahikainen, Anthony H. Keeble, Lisa Schimanski, Saira Hussain, Ruth Harvey, Jack W.P. Hayes, Jane. C. Edwards, Rebecca K. McLean, Veronica Martini, Miriam Pedrera, Nazia Thakur, Carina Conceicao, Isabelle Dietrich, Holly Shelton, Anna Ludi, Ginette Wilsden, Clare Browning, Adrian K. Zagrajek, Dagmara Bialy, Sushant Bhat, Phoebe Stevenson-Leggett, Philippa Hollinghurst, Matthew Tully, Katy Moffat, Chris Chiu, Ryan Waters, Ashley Gray, Mehreen Azhar, Valerie Mioulet, Joseph Newman, Amin S. Asfor, Alison Burman, Sylvia Crossley, John A. Hammond, Elma Tchilian, Bryan Charleston, Dalan Bailey, Tobias J. Tuthill, Simon P. Graham, Tomas Malinauskas, Jiandong Huo, Julia A. Tree, Karen R. Buttigieg, Raymond J. Owens, Miles W. Caroll, Rodney S. Daniels, John W. McCauley, Kuan-Ying A. Huang, Mark Howarth, Alain R. Townsend

## Abstract

There is dire need for an effective and affordable vaccine against SARS-CoV-2 to tackle the ongoing pandemic. In this study, we describe a modular virus-like particle vaccine candidate displaying the SARS-CoV-2 spike glycoprotein receptor-binding domain (RBD) using SpyTag/SpyCatcher technology (RBD-SpyVLP). Low doses of RBD-SpyVLP in a prime-boost regimen induced a strong neutralising antibody response in mice and pigs that was superior to convalescent human sera. We evaluated antibody quality using ACE2 blocking and neutralisation of cell infection by pseudovirus or wild-type SARS-CoV-2. Using competition assays with a monoclonal antibody panel, we showed that RBD-SpyVLP induced a polyclonal antibody response that recognised all key epitopes on the RBD, reducing the likelihood of selecting neutralisation-escape mutants. The induction of potent and polyclonal antibody responses by RBD-SpyVLP provides strong potential to address clinical and logistic challenges of the COVID-19 pandemic. Moreover, RBD-SpyVLP is highly resilient, thermostable and can be lyophilised without losing immunogenicity, to facilitate global distribution and reduce cold-chain dependence.

## INTRODUCTION

Coronavirus disease 2019 (COVID-19), caused by a novel coronavirus named severe acute respiratory syndrome coronavirus 2 (SARS-CoV-2), was first reported in Wuhan, China in December 2019 ^1^. Since then COVID-19 has spread across the world and was declared a pandemic by the World Health Organisation (WHO) in March 2020. As of August 2020, there have been over 20 million confirmed COVID-19 cases worldwide and around 800,000 deaths ^2^. There are no vaccines or effective treatments for COVID-19 to date; however, as of August 2020, there are 25 vaccine candidates in clinical evaluation and around 140 are in pre-clinical testing ^3^. Vaccine candidates in current clinical evaluation include inactivated, viral vector (replicating and non-replicating), protein subunit, nucleic acid (DNA and RNA) and virus-like particle (VLP) vaccines with the majority of them focusing on using the full-length SARS-CoV-2 spike glycoprotein (S) as an immunogen.

SARS-CoV-2 is an enveloped virus carrying a single-stranded positive-sense RNA genome (∼30 kb), belonging to the genus *Betacoronavirus* from the *Coronaviridae* family ^4^. The virus RNA encodes four structural proteins including spike (S), envelope (E), membrane (M), and nucleocapsid (N) proteins, 16 non-structural proteins, and nine accessory proteins ^5^. The S glycoprotein consists of an ectodomain (that can be processed into S1 and S2 subunits), a transmembrane domain, and an intracellular domain ^6^. Similar to the Severe Acute Respiratory Syndrome Coronavirus (SARS-CoV), SARS-CoV-2 binds the human angiotensin-converting enzyme 2 (ACE2) via the receptor-binding domain (RBD) within the S1 subunit to facilitate entry into host cells, followed by membrane fusion mediated by the S2 subunit ^7-9^

Of the many vaccine platforms, protein subunit vaccines generally have good safety profiles and their production is rapid and easily scalable ^10^. Recombinant RBD proteins of SARS-CoV and MERS-CoV have been shown to be immunogenic and induce protective neutralising antibodies in animal models and are therefore considered promising vaccine candidates (reviewed ^11, 12^). RBD from SARS-CoV-has recently been confirmed to be inducing neutralising antibodies ^13, 14^. Recently published studies, including one from our group, found that the majority of the potent neutralising antibodies isolated from SARS-CoV-2-infected patients bound to the RBD ^15, 16^. We therefore chose to study the immunogenicity of RBD. To improve immunogenicity, we conjugated the RBD onto a virus-like particle (VLP). VLP display of protein antigen has been shown to further enhance immunogenicity by facilitating antigen drainage to lymph nodes, enhancing uptake by antigen-presenting cells and increasing B cell receptor crosslinking ^10, 17^. Moreover, we recently showed that influenza antigens (haemagglutinin (HA) or neuraminidase (NA)) displayed on the VLPs (same VLP used in this study) were highly immunogenic at a low dose (0.1 µg) in mice ^18^.

In the present study, we used the SpyTag/SpyCatcher technology for assembly of SARS-CoV-2 RBD on the mi3 VLP, via the formation of an intermolecular isopeptide bond between the RBD and the VLP, as a potential SARS-CoV-2 vaccine ^19, 20^ (Figure 1A). SpyTag-mediated VLP decoration has been successfully used for the display of diverse antigens from, e.g. *Plasmodium spp*., influenza A virus, HIV and cancer cells (PD-1L) ^18, 19, 21, 22^. The RBD-SpyVLP vaccine candidate is highly immunogenic in mice and pigs, inducing robust SARS-CoV-2 neutralising antibody responses. The results of our study demonstrate the potential of the RBD-SpyVLPs as an effective and affordable vaccine for COVID-19 with RBD-SpyVLP being resilient, retaining stability and immunogenicity post-lyophilisation, which will greatly facilitate distribution for vaccination by eliminating cold-chain dependence.

**Figure 1.**
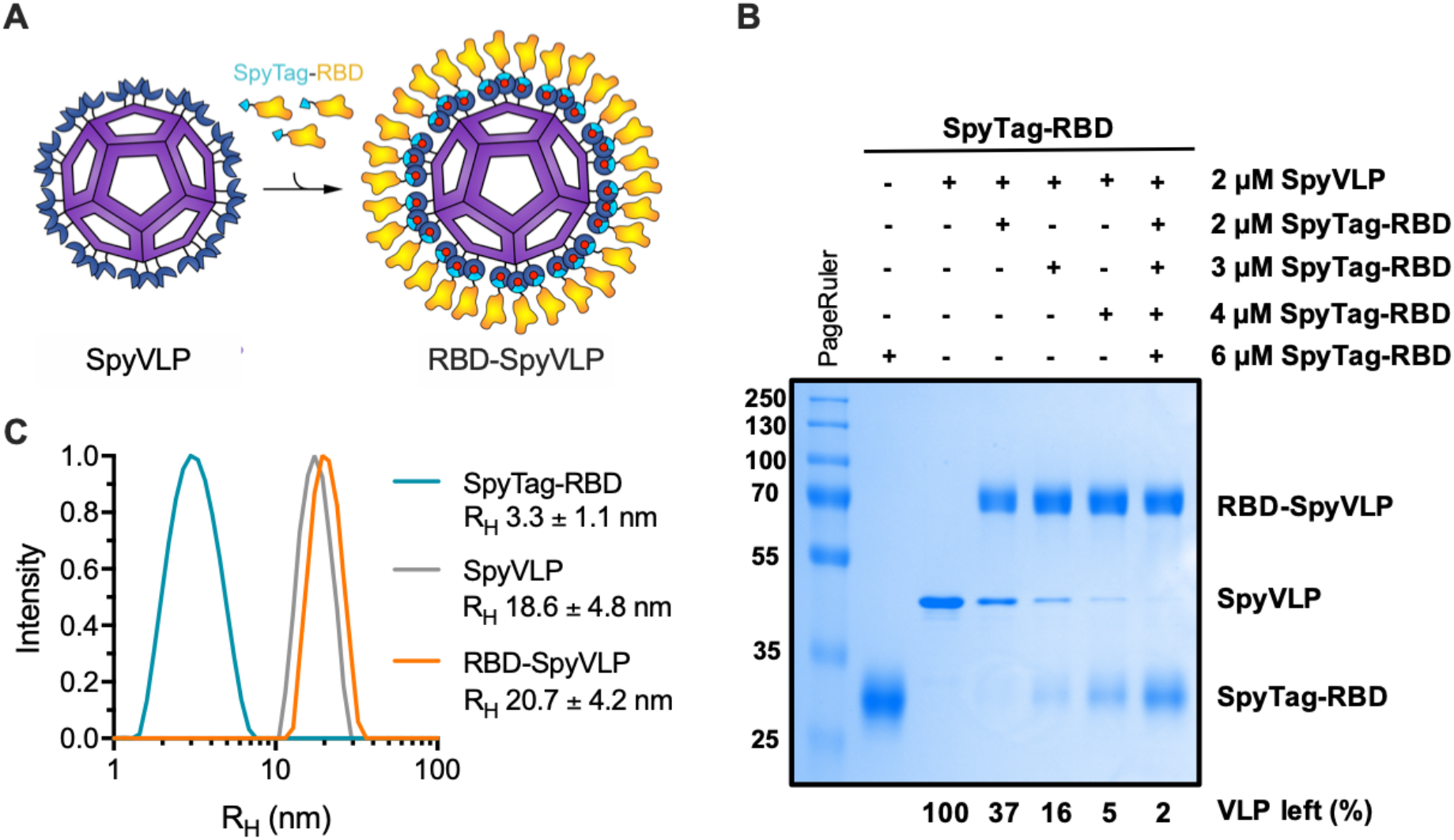
SpyTag-RBD can be efficiently conjugated to SpyCatcher003-mi3 VLP. (A) Schematic diagram of the RBD-SpyVLP vaccine candidate, consisting of SpyCatcher003-VLP conjugated with SpyTag-RBD. The isopeptide bonds formed spontaneously between SpyTag and SpyCatcher are indicated with red dots. (B) Conjugation of SpyCatcher003-mi3 with SpyTag-RBD at various ratios. Reactions were performed at 4 °C overnight and analysed using SDS-PAGE with Coomassie staining and densitometry, with the percentage of unreacted VLP shown. (C) Dynamic light scattering (DLS) characterisation of SpyTag-RBD, SpyVLP, and conjugated RBD-SpyVLP (n=3, values shown as mean±SD). R_H_= hydrodynamic radius.

## RESULTS

### 1. RBD can be efficiently displayed on the mi3 VLP via SpyTag/SpyCatcher

To create the RBD-SpyVLP vaccine candidate, the SpyTag (AHIVMVDAYKPTK) coding sequence was fused between the signal sequence from influenza H7 HA and the N-terminus of the RBD (amino acid 340-538, NITN…GPKK) (A/HongKong/125/2017) (SpyTag-RBD) (see Figure S1 for the full sequence) and the glycoprotein was expressed in mammalian cells (Expi293) prior to purification using Spy&Go affinity chromatography ^23^. The purified SpyTag-RBD was then conjugated to the SpyCatcher003-mi3 VLP ^18, 20^ to generate the SpyTag-RBD:SpyCatcher003-mi3 (RBD-SpyVLP) immunogen (Figure 1A). SpyCatcher003 is a variant of SpyCatcher which was engineered for accelerated reaction with SpyTag ^24^. mi3 is a computationally engineered dodecahedron based on an aldolase from a thermophilic bacterium ^20, 25^. The SpyTag-RBD can be efficiently conjugated to the SpyCatcher-mi3 VLP, with 93% display efficiency reached after 16 h (Figure 1B). This corresponds to an average of 56 RBDs per VLP. We saw no sign of VLP aggregation following coupling and the RBD-SpyVLP is homogeneous, as shown by a uniform peak of the hydrodynamic radius (R_H_) at 20.7 ± 4.2 nm in dynamic light scattering (DLS) (Figure 1C). For immunisation, we chose a conjugation ratio that leaves minimal free RBD (1:1 molar ratio), which corresponds to ∼64% display efficiency or around 38 RBD per VLP (Figure 1B).

### 2. RBD-SpyVLP is reactive to monoclonal antibodies isolated from recovered patients and is resilient

To confirm the antigenicity of RBD-SpyVLP, we performed a series of binding assays. Binding to RBD-SpyVLP was tested using a panel of novel monoclonal antibodies (mAbs) some of which are strongly neutralising, from COVID-19-infected donors ^26^, that bind to at least three independent epitopes on the RBD. We included the published conformation-specific mAb (CR3022) ^27^, a nanobody-Fc fusion VHH72-Fc ^28^, and a dimeric human ACE2-Fc ^29^, for which there are published structures. All tested mAbs and ACE2-Fc bound strongly to the RBD-SpyVLP (Figure 2A), showing that a broad range of epitopes on the RBD-SpyVLP are exposed and correctly folded. An anti-influenza neuraminidase mAb (Flu mAb), used as a negative control, showed no binding to RBD-SpyVLP, confirming the specificity of the assay (Figure 2A).

**Figure 2.**
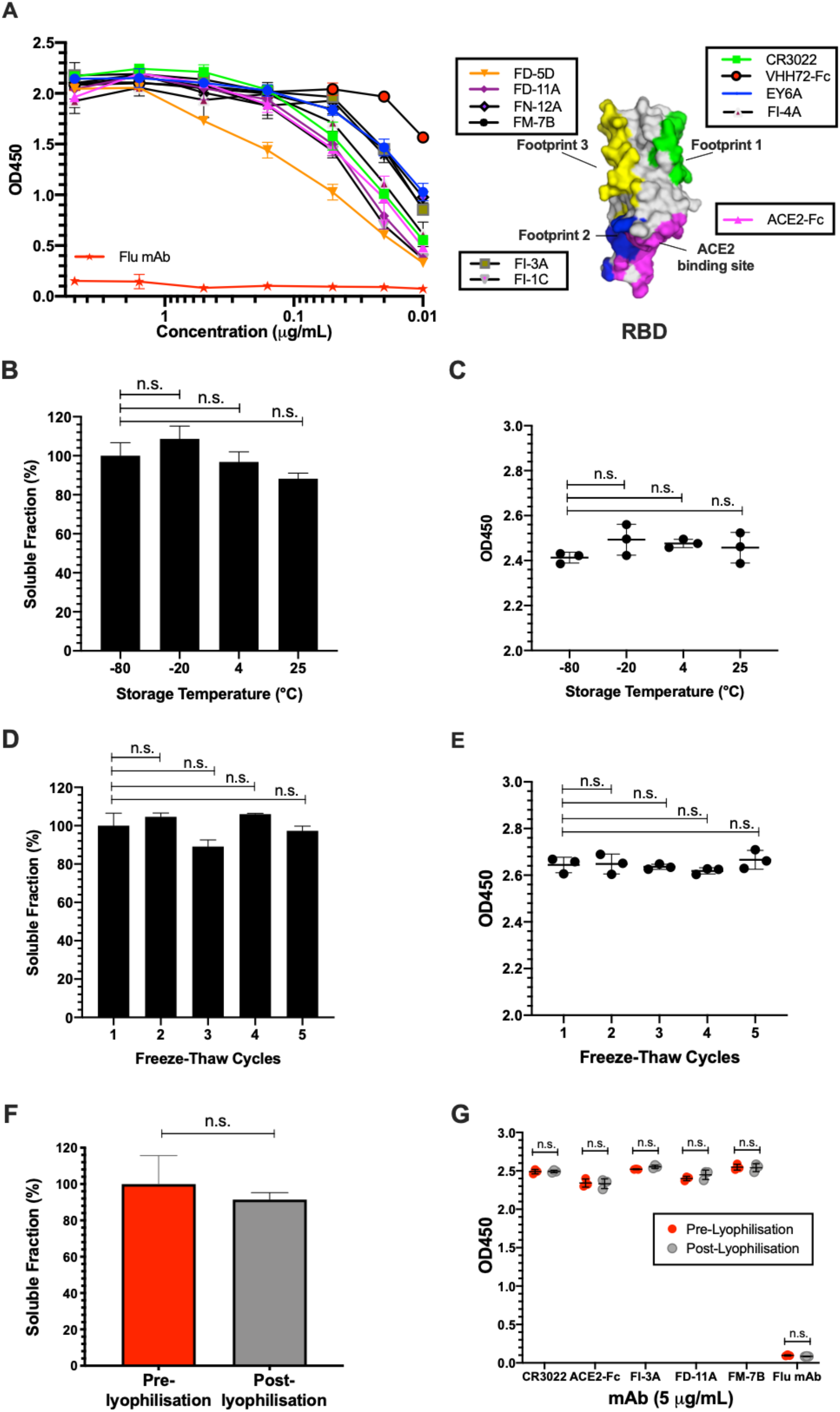
RBD-SpyVLPs are reactive to SARS-CoV-2 binders and thermostable and resilient. (A) Binding of RBD-SpyVLP to a panel of monoclonal antibodies isolated from COVID-19 recovered patients that target independent epitopes on the RBD determined using competitive ELISA (Huang et al., 2020). The boxed antibodies form groups that compete with each other and with antibodies or nanobodies with structurally defined footprints: CR3022 (PDB 6W41) (footprint 1), H11-D4-Fc (footprint 2) (PDB 6YZ5), S309 (footprint 3) (PDB 6WPT) and the ACE2 binding site (PDB 6M0J). Each point represents the mean of duplicate readings and error bars represent ± 1 SD. The diagram of the RBD, created in PyMOL, shows the ACE2 binding site and the three binding footprints highlighted. (B) Solubility and (C) immunoreactivity of RBD-SpyVLP after storage for two weeks at various temperatures, determined using SDS-PAGE and densitometry or ELISA (10 µg/mL CR3022 mAb). (D) Solubility and (E) immunoreactivity of RBD-VLP after freeze−thaw determined using SDS-PAGE and ELISA (10 µg/mL CR3022 mAb) after one to five cycles of freeze−thawing. (F) RBD-SpyVLP soluble fraction, before and after lyophilisation reconstituted in the same buffer volume and (G) immunoreactivity determined using ELISA with ACE2-Fc and mAbs that target non-overlapping epitopes on the RBD. Error bars in B, E & F represent group mean (n=3). Error bars in C, E & G represent mean ± 1 SD (n= 3). Statistical difference in B to E was determined using Kruskal-Wallis test followed by Dunn’s multiple comparison test. Statistical difference in F & G was determined using Mann-Whitney U test. n.s. = not significant.

We then tested the stability of RBD-SpyVLP, to determine its resilience and likely sensitivity to failures in the cold-chain ^30^. The unconjugated SpyCatcher003-mi3 VLP had previously been shown to be highly thermostable as a platform for antigen display ^18^. For conjugated RBD-SpyVLP, we tested its solubility following storage for two weeks at −80, −20, 4 or 25 °C in Tris Buffered Saline (TBS). We then centrifuged out any aggregates and analysed soluble protein by SDS-PAGE with Coomassie staining. We found no significant change in the soluble fraction following storage at 4 °C (n=3, Kruskal-Wallis and Dunn’s post-hoc test, p>0.05), with only a 12% decrease after storage for two weeks at 25 °C (Figure 2B) and no degradation was observed at 25 °C (Figure S2A). We further analysed the integrity of the sample with ELISA against the conformation-dependent CR3022 mAb and observed no loss of antigenicity under these storage conditions (Figure 2C). We next assessed the resilience of RBD-SpyVLP to freezing, challenging RBD-SpyVLP with multiple rounds of freeze-thaw. Even after five rounds of freeze-thaw, there was no significant loss of soluble RBD-SpyVLP (Figure 2D) or CR3022 recognition (Figure 2E) (n=3, Kruskal-Wallis and Dunn’s post-hoc test, p>0.05) and no degradation was observed (Figure S2B). After reconstitution following lyophilisation, we saw a minimal change in soluble protein for RBD-SpyVLP (91.5±3.8% of the initial value (mean±SD) (Figure 2F), which was not statistically significant (n=3, Mann-Whitney U test, p>0.05). There was also no difference in terms of binding of RBD-SpyVLP to a panel of mAbs or ACE2-Fc recognising non-overlapping footprints on the RBD (Figure 2G). Overall, RBD-SpyVLP showed a high level of resilience.

### 3. RBD-SpyVLP induces a strong ACE2-blocking and neutralising antibody response in mouse models

We first evaluated the immunogenicity of RBD-SpyVLP in mouse models. C57BL/6 mice (n=6) were immunised intramuscularly (IM) with purified RBD alone (0.1 µg or 0.5 µg), RBD-SpyVLP (0.1 µg or 0.5 µg equivalents of the RBD component) or VLP alone, all adjuvanted with AddaVax. AddaVax is a squalene-based oil-in-water nano-emulsion adjuvant, a pre-clinical equivalent to the licensed MF59 adjuvant ^31^. Mice were then boosted with the same dose of immunogen two weeks later and sera were collected at three weeks post-boost. Both the 0.1 µg and 0.5 µg RBD-only groups showed levels of antibody against RBD or spike glycoprotein only slightly above background, detected using ELISA (serum reciprocal endpoint titre (EPT): 1:94 and 1:68, respectively), and showed no difference compared to the VLP-only group (Figure 3A and B). Mice immunised with 0.1 µg and 0.5 µg RBD-SpyVLP groups showed high levels of antibody to RBD (EPT: 0.1 µg: 1:16,117, p<0.001 and 0.5 µg: 1:7,300, p<0.01) (Figure 3A) and to spike (EPT: 0.1 µg: 1:1,647, p<0.05 and 0.5 µg: 1:1,212) compared to the VLP group (Figure 3B).

**Figure 3.**
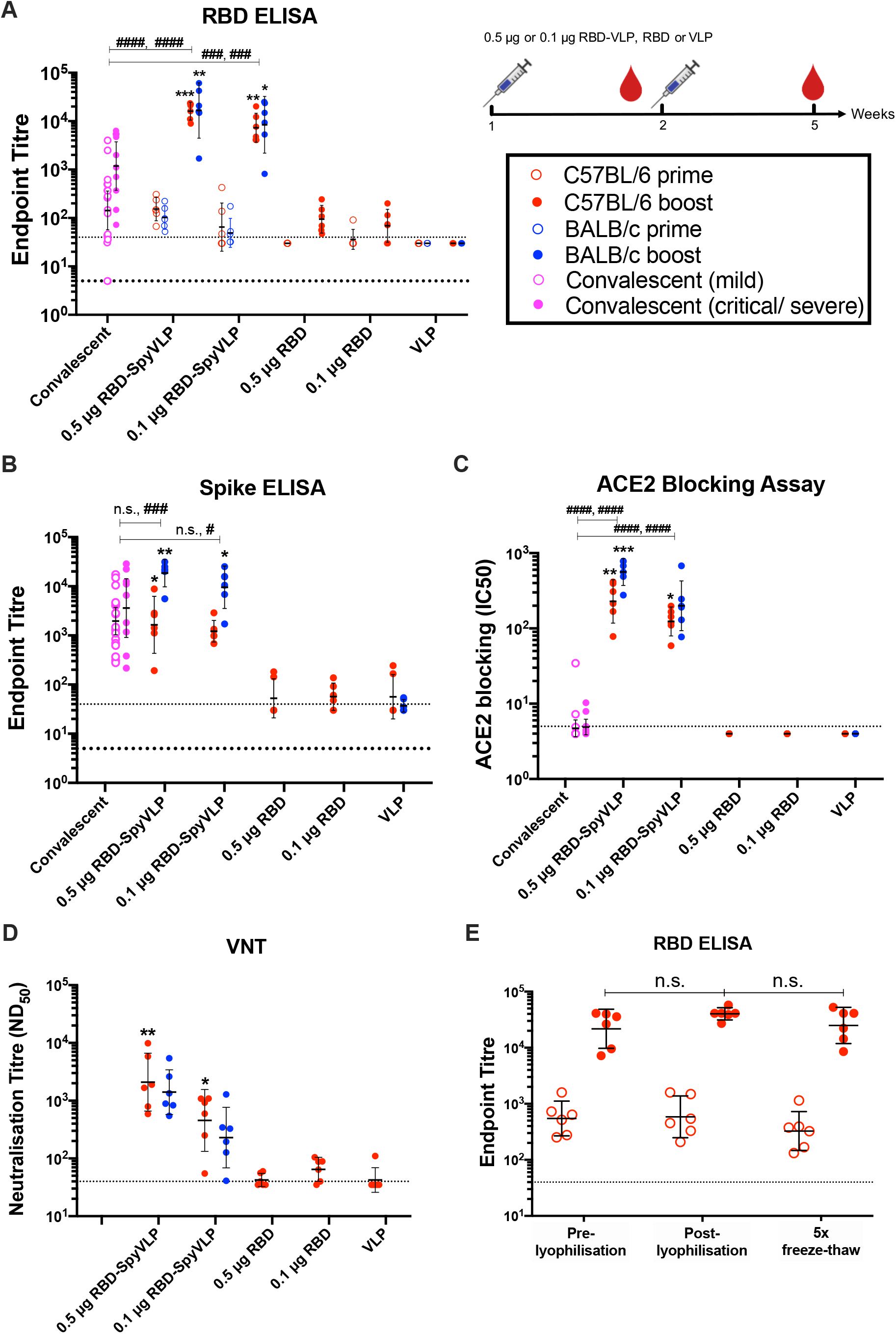
RBD-SpyVLPs induce strong antibody responses in mice that are comparable to the responses in recovered patients. C57BL/6 (red) or BALB/c (blue) mice (n=6 in each group) were dosed twice IM, two weeks apart with 0.1 µg or 0.5 µg purified RBD, RBD-SpyVLP or VLP alone with AddaVax adjuvant added to all. Sera were harvested at two weeks after the first dose (open circles) and at three weeks after the second dose (closed circles). Sera were analysed in (A) ELISA against RBD ELISA against full-length spike glycoprotein, (C) in an ACE2 competition assay, and (D) in virus neutralisation assays (VNT) against wild-type SARS-CoV-2 virus. (E) Antibody response analysed by RBD ELISA for mice dosed twice with 0.5 µg RBD-SpyVLP (pre-lyophilised, post-lyophilised or freeze-thawed 5 times (5x FT). Data are presented as the group geometric means ± 95% confidence intervals. COVID-19 convalescent plasma from humans with mild (open mauve circles) or critical/severe (closed mauve circles) disease were included for comparison. *p<0.05, ** p<0.01, ***p<0.001 determined by Kruskal-Wallis test followed by Dunn’s multiple comparison test. ^#^p<0.05, ^###^p<0.001, ^####^p<0.0001 determined by Mann-Whitney U test to compare against convalescent human plasma. Dotted lines represent the lowest mouse sera dilutions tested. Bold dotted line represents the lowest human sera dilution tested.

We then tested RBD-SpyVLP in a second mouse strain (BALB/c) with the same dosage regimen to confirm the immunogenicity (Figure 3A and B). In BALB/c, the 0.1 µg and 0.5 µg RBD-SpyVLP groups showed higher levels of RBD-specific antibody (EPT: 0.1 µg: 1:8,406, p<0.01 and 0.5 µg: 1:16,636, p<0.05) (Figure 3A) and spike-glycoprotein specific antibody (EPT: 0.1 µg: 1:9,574, p<0.001 and 0.5 µg: 1:18,556, p<0.05) (Figure 3B) compared to the VLP group.

The ability of immunised mouse sera to block recombinant soluble ACE2 binding to immobilised RBD was then assessed. Sera from the 0.1 µg and 0.5 µg RBD-SpyVLP immunised C57BL/6 mice had significantly higher ACE2 blocking (IC_50_: 0.1 µg: 1:132, p<0.05 and 0.5 µg: 1:253; p<0.05) activity compared to the VLP-immunised mice (Figure 3C). C57BL/6 mice immunised with RBD-only showed no detectable ACE2 blocking activity, consistent with the ELISA results (Figure 3A & B). Similarly, both 0.1 µg and 0.5 µg RBD-SpyVLP-immunised BALB/c mice had significantly higher ACE2 blocking (IC_50_: 0.1 µg:1:200, p<0.001 and 0.5 µg: 1:560, respectively) compared to the VLP immunised group (Figure 3C). In both mouse strains, 0.1 µg and 0.5 µg RBD-SpyVLP immunised groups had higher antibody titres against RBD (around 5-10-fold; C57BL/6, p<0.001 and p<0.0001; BALB/c, p<0.001 and p<0.0001) and ACE2 blocking (around 10-50-fold; C57BL/6, p<0.0001 and p<0.0001; BALB/c, p<0.0001 and p<0.0001) compared to plasma donated by patients convalescing from COVID-19 disease (n=28) (Figure 3B and C). Sera from BALB/c mice immunised with 0.1 µg and 0.5 µg RBD-SpyVLP had comparable spike glycoprotein-specific antibody responses compared to convalescent humans whereas sera from C57BL/6 mice immunised with either dose had higher spike gluycoprotein-specific antibody responses compared to convalescent humans (around 2-fold, 0.1 µg, p<0.05 and 0.5 µg, p<0.001) (Figure 3B).

The antibody response in mice was assessed for neutralisation potency using a live SARS-CoV-2 virus (hCoV-19/England/02/2020, EPI_ISL407073) neutralisation assay (VNT) based on virus plaque reduction. Sera from C57BL/6 mice immunised with either dose of unconjugated RBD showed low level neutralising titres (ND_50_ 1:59 and 1:32) compared to the RBD-SpyVLP group, again consistent with ELISA and ACE2 blocking activity (Figure 3D). Both 0.1 µg and 0.5 µg RBD-SpyVLP immunised groups exhibited neutralising titres in both C57BL/6 (ND_50_: 1:450 to 1: 2,095) and BALB/c mice (ND_50_: 1:230 to 1:1,405) (Figure 3D). Consistent neutralising activity in sera from C57BL/6 mice was found using a VNT in an independent laboratory with the hCoV-19/VIC01/2020 isolate (GenBank MT007544) (Table S1).

We tested the immunogenicity of RBD-SpyVLP post-lyophilisation and after five freeze-thaw cycles. C57BL/6 mice (n=6) were immunised IM with 0.5 µg RBD-SpyVLP (pre-lyophilised, post-lyophilised or 5× freeze-thaw) and sera were harvested post prime and post boost and tested for binding in a RBD ELISA. There was no difference between the immune responses in all three groups tested (Figure 3E) (Kruskal-Wallis, followed by Dunn’s post-hoc test, p>0.05), showing that lyophilisation and freeze-thaw had not compromised the immunogenicity of the RBD-SpyVLP.

### 4. RBD-SpyVLP is highly immunogenic and induces strong neutralising antibody response in pigs

We tested RBD-SpyVLP for its immunogenicity in pigs as a large, genetically outbred animal model. Pigs have previously been reported to be a reliable model to study vaccines for use in humans because of their highly similar physiologies and immune systems ^32, 33^. The pig model has been used recently to test an adenovirus vector vaccine candidate against SARS-CoV-2 (ChAdOx1 nCoV19) ^34^. We immunised pigs (n=3) with a dose of RBD-SpyVLP that we intend to use in humans (5 µg) or with a 10-fold higher dose (50 µg) to study dose-response. A third group (n=3) receiving 100 µg of purified trimeric spike glycoprotein (spike) was included as a control. Pigs were immunised IM twice, 28-days apart, with 5 µg or 50 µg RBD-SpyVLP or spike, all adjuvanted with AddaVax.

As early as day 7, pigs immunised with 50 µg RBD-SpyVLP showed a detectable anti-RBD antibody response, whereas no antibody response was detected at day 7 for 5 µg RBD-SpyVLP or 100 µg spike groups (Figure 4A) (p<0.05, Kruskal-Wallis test). The antibody response in all three groups increased gradually to day 14 when the response reached a plateau. The group receiving 50 µg RBD-SpyVLP group showed a trend of slightly higher antibody response than the other two groups until the day of boost (day 28) (Figure 4A). The antibody response in all three groups increased by around 100-fold one week after boosting and remained high until at least day 56. There was no dose-response difference between the 5 µg and 50 µg RBD-SpyVLP groups (p>0.5, Kruskal-Wallis test) for day 56 data (Figure 4A).

**Figure 4.**
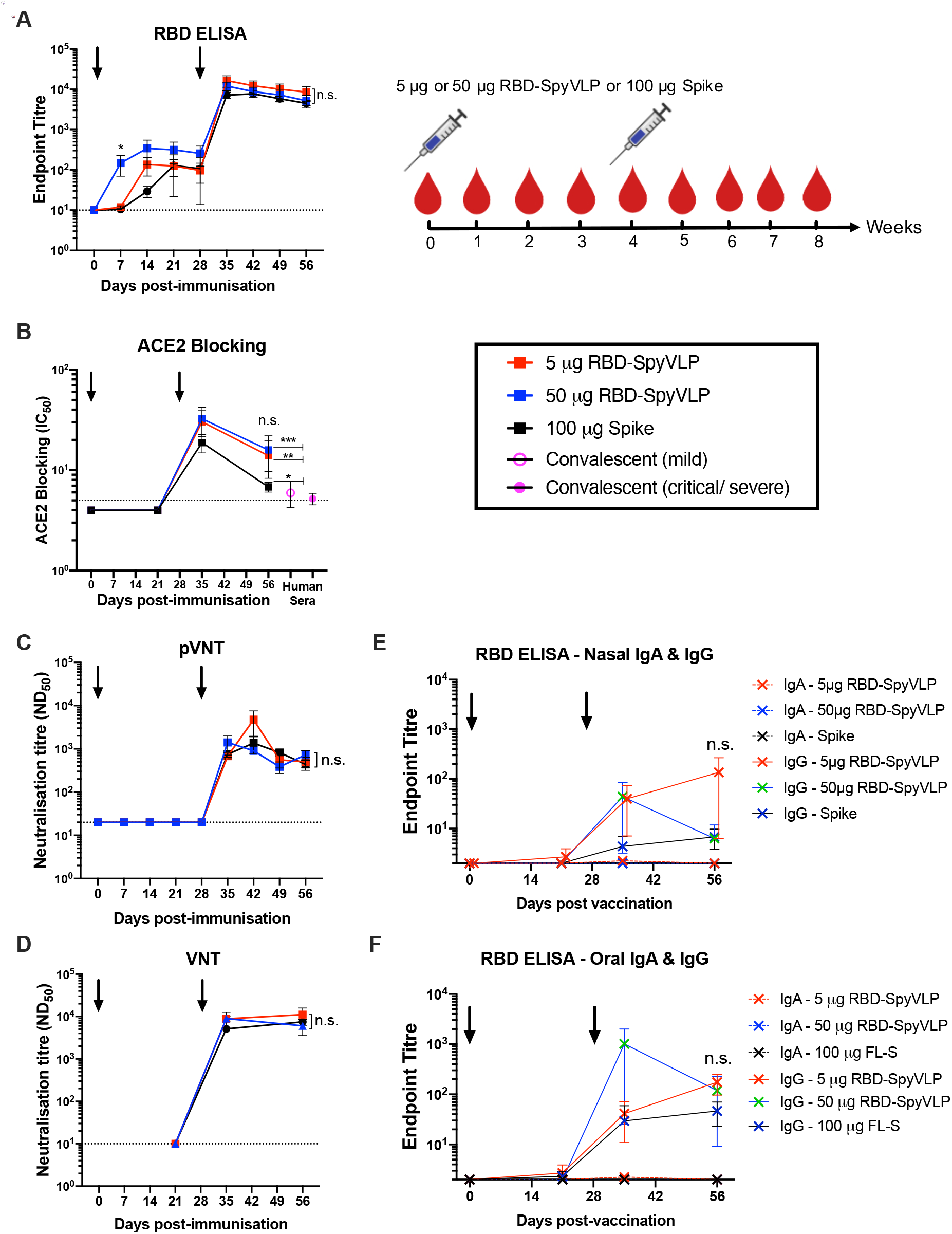
RBD-SpyVLPs induce persistent and strong neutralising antibody responses in pigs. Pigs (n=3 in each group) were dosed twice (prime and boost) IM, a month apart with 5 µg or 50 µg RBD-VLP or 100 µg of spike glycoprotein, all adjuvanted with AddaVax and sera were harvested at indicated time-points and analysed in (A) ELISA against RBD, (B) in an ACE2 competition assay, (C) in pseudovirus neutralisation assay (pVNT), and (D) wild-type virus neutralisation assay (VNT). Antibody levels in (E) nasal and (F) oral swabs were measured at indicated time points using RBD ELISA. Data are presented as the group mean ± 1 SEM. Black arrows indicate when the vaccines were administered. Kruskal-Wallis test was used to compare data on day 56 in B-E, n.s.= not significant. *p<0.05, **p<0.01 and ***p<0.001 on day 56 data compared to convalescent human plasma determined by Mann-Whitney U Test. Dotted lines represent the lowest dilutions of sera tested in the assays.

ACE2 blocking activity in the pig serum was measured at day 21, day 35 and day 56. No ACE2 blocking was detected at day 21, prior to the booster dose. ACE2 blocking was detected post-boost at day 35 (IC_50_ 1:10 to 1:50) and reduced at day 56 (IC_50_ 1:5 to 1:25) in all three groups (Figure 4B). ACE2 blocking activities in all three groups were significantly higher than for sera from convalescent humans (5 µg RBD-SpyVLP, p<0.01; 50 µg RBD-SpyVLP, p<0.001; 100 µg, p<0.05), by around ∼3-6-fold (p<0.05 and p<0.001, respectively, Kruskal-Wallis test on day 56 data).

Neutralising antibody responses in the pigs were assessed against both SARS-CoV-2 pseudotyped lentivirus and live SARS-CoV-2 virus. All three groups exhibited similar neutralisation titres against pseudovirus (ND_50_: ∼1:500 to 1:4,000) after boost which remained until at least day 56, with no difference between the three groups (p>0.05, Kruskal-Wallis test on day 56) (Figure 4C). Similarly, no difference in neutralising activity was detected in all three groups against live virus at day 21 (Figure 4D). Neutralisation was detected on both day 35 and 56 in all groups, with ND_50_ levels between 1:6,000 to 1:11,000 on day 56 and no significant difference between the three groups (p>0.05, Kruskal-Wallis test on day 56 data) (Figure 4D). Nasal and oral secretions were also tested for the presence of neutralising antibodies: here RBD-specific IgG, but notably not IgA, was detected in all three groups with no significant difference between groups post-boost for both nasal and oral secretions (p>0.05, Kruskal-Wallis followed by Dunn’s Post-hoc) (Figure 4E & F).

We assessed the level of RBD-specific B cells in peripheral blood. The predominance of an IgG response following the booster immunisation was confirmed in all three groups by assessment of RBD tetramer labelling of IgM, IgG and IgA B cells in peripheral blood and IgG ELISpot assay (Figure S3A & B). In each case, RBD-specific IgG B cells peaked in peripheral blood shortly after the boost (Figure S3A). We also performed longitudinal analysis of CD4^+^ and CD8^+^ T cell responses following immunisation. Intracellular cytokine staining of S-peptide-stimulated peripheral blood mononuclear cells (PBMCs) demonstrated a T cell IFN-γ response, slightly larger in the CD4^+^ than CD8^+^ T cell pool, which peaked 7 days after prime and boost immunisations (Figure S3C).

### 5. RBD-SpyVLP induces polyclonal antibody responses against the RBD in mice and pigs

A concern regarding RBD-based vaccines is whether the immune response will be focussed to a single site on the antigen, because of the relatively small size of the RBD, potentially leading to a narrow response that would be sensitive to immune escape ^35, 36^. A recent report showed that passive immunisation with a single mAb led to escape mutants of SARS-CoV-2, whereas a cocktail of two neutralising antibodies to independent epitopes prevented emergence of neutralisation-escape mutations, demonstrating the importance of a polyclonal antibody response ^37^. We assessed sera from mice and pigs immunised with RBD-SpyVLP for antibody responses that target multiple RBD epitopes using a competition ELISA against four different mAbs: FI-3A, FD-11A, EY6A and S309, that target three non-overlapping epitopes on the RBD with FD-11A and S309 showing overlap as defined by competition ELISA (see diagram in Figure 2A) ^26, 38, 39^,. BALB/c mice immunised with RBD-SpyVLP showed competition against all four mAbs tested (Figure 5A). BALB/c mice immunised with VLP-only showed no competition (p<0.05, Mann Whitney U-Test) (Figure 5A). A similar pattern was observed in the C57BL/6 mice (Figure S5). Comparison of preimmune and day 42 post-RBD-SpyVLP-immunisation sera from pigs showed a trend of partial competition against all four mAb, but this was not statistically significant because of inter-animal variation (Figure 5B). When responses of individual pigs were compared to their pre-immune sera 2 out of 3 animals in each dosing group showed significant competition for the antibodies FI-3A and S309 that defined independent neutralising epitopes compared to their preimmune sera (Figure S5). These results show that RBD-SpyVLP does not have an immunodominant epitope and does not induce a highly focussed antibody response, making it a vaccine candidate that is likely to resist the generation of neutralisation-escape mutants.

**Figure 5.**
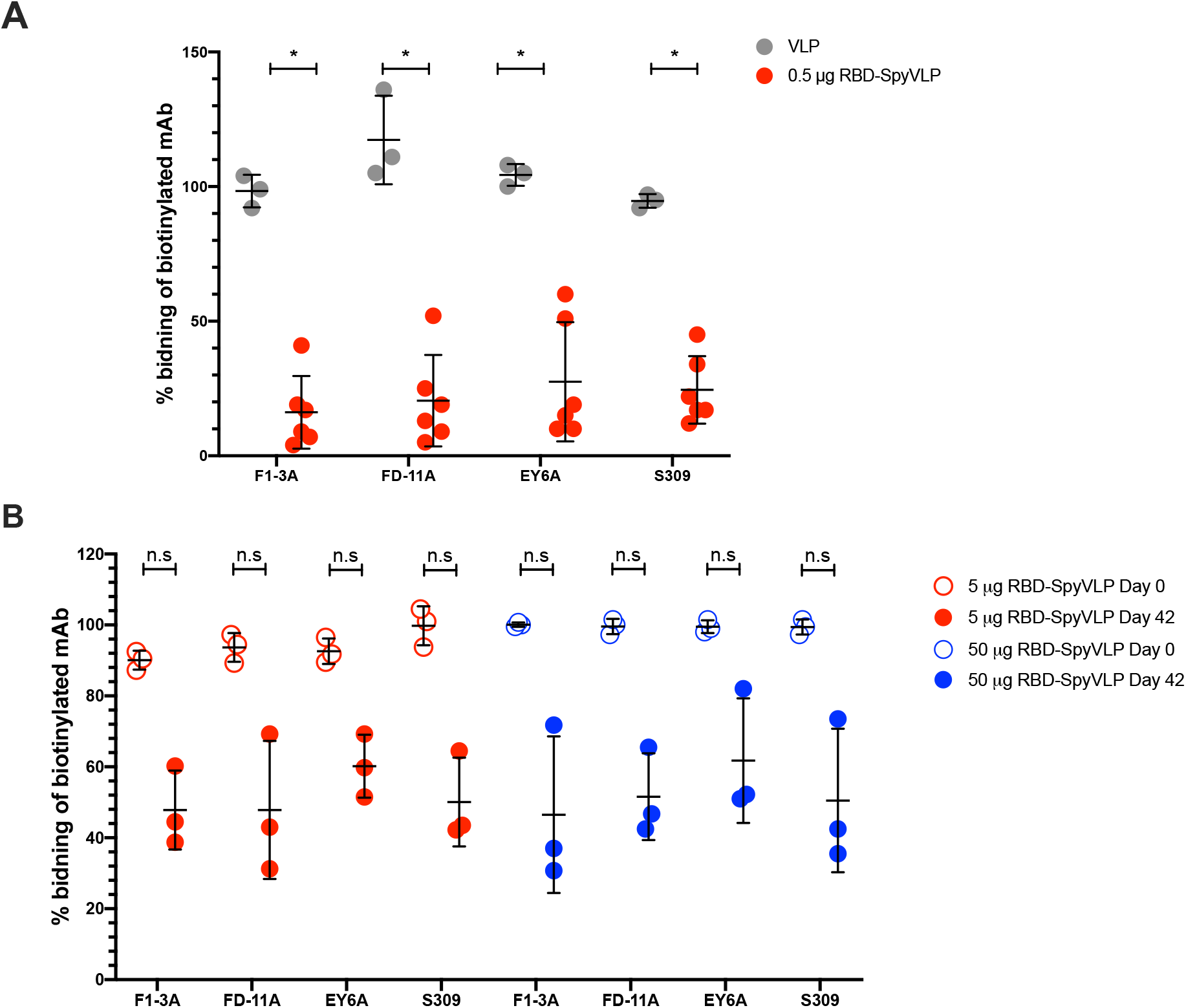
RBD-SpyVLP immunisation of mice and pigs elicits polyclonal antibody responses that targets all key epitopes on the RBD. Competition ELISA of four human mAbs that target three different key epitopes on the RBD with post-boost BALB/c sera (0.5 µg RBD-SpyVLP) (A). Competition ELISA of four human mAbs that target three different key footprints on the RBD with post-boost pig sera (5 µg or 50 µg RBD-SpyVLP) compared to preimmune sera (day 0) (B). Each point in A represents an average of duplicate readings of a serum sample from one animal three weeks post boost tested at 1:20 dilution. Each point in B represents an average of quadruplicate readings of a serum sample from one animal on day 42 tested at 1:20 dilution Data are presented as group means ± 1 SD. ** p<0.01, determined by Mann-Whitney U test. n.s. = not significant.

## DISCUSSION

In the current study we investigated an RBD-based VLP vaccine candidate for COVID-19 based on SpyTag/SpyCatcher technology which was used to assemble RBDs into the mi3 VLP via the formation of an irreversible isopeptide bond ^19^. We showed RBD-SpyVLPs to be strongly immunogenic in mice and pigs, inducing high titre neutralising antibody responses against wild type SARS-CoV-2 virus. This study confirms that the RBD is the key immunogenic domain for eliciting neutralising monoclonal antibodies against SARS-CoV-2, in line with studies showing that highly neutralising antibodies isolated from convalescent patients bind to the RBD ^16, 26, 38, 40-46^. We showed that RBD-SpyVLPs are recognised by a panel of mAbs isolated from convalescent patients ^26^ binding to various epitopes on the RBD (Figure 2A). This distributed reactivity shows that all of the epitopes that could potentially induce protective antibodies to RBD are present in RBD-SpyVLPs.

We detected negligible antibody responses in mice vaccinated with equivalent doses (0.1 µg or 0.5 µg) of purified RBD alone, but strong responses to the RBD when displayed on the VLP (Figure 3). Previous studies showed that RBD from SARS-CoV and SARS-COV-2 can induce neutralising antibodies in animal models but typically after administration of much higher doses (e.g. ∼50 to 100 µg) and with frequent dosing ^13, 14, 47^. On the other hand, we showed that high titre neutralising antibody responses can be detected in two strains of mice immunised with relatively low doses of RBD-SpyVLP (to ND_50_ ∼500 to 2,000). These results confirm the enhanced immunogenicity of RBD when displayed on SpyVLPs. Sera from mice immunised with both 0.1 µg or 0.5 µg of RBD-SpyVLP exhibited high levels of antibody against SARS-CoV-2 RBD and full-length spike glycoprotein and ACE2 blocking activity (Figure 3B, C and D). All of these responses were higher than the levels found in plasma from convalescent humans. Together, these observations suggest that RBD-SpyVLP vaccination could potentially elicit protective antibody responses against SARS-CoV-2 in humans.

RBD-SpyVLP vaccination also induces high titre neutralising antibody responses in pigs (ND_50_ ∼1:11,000) with a dose that we aim to use for subsequent human trials (5 µg) (Figure 4D). At a dose around 2-fold less (based on molar ratio), 5 µg of RBD-SpyVLP induced similar neutralisation titres compared to 100 µg of spike glycoprotein, showing the excellent immunogenicity of RBD-SpyVLP vaccination. Transudate RBD-specific IgG from serum can be detected in the oral and nasal cavity (Figure 4E & F). Surprisingly, no increase in antibody titre was observed in pigs that received a higher dose of antigen (50 µg RBD-SpyVLP). There was a trend that the 50 µg RBD-SpyVLP group generated a more rapid and higher response post-prime but the antibody response between the 5 µg and 50 µg RBD-SpyVLP groups were identical post-boost. This suggests a threshold effect.

Since RBD-SpyVLPs induce antibody responses that target multiple epitopes on the RBD the chance of selecting neutralisation-escape mutants should be greatly reduced. Circulating SARS-CoV-2 stains are constantly mutating and the likelihood of persistence of the virus in the human population is high ^48, 49^.

We observed differences in the levels of ACE2 blocking in sera from immunised mice and pigs (Figure 3C and 4B). Sera taken from mice immunised with RBD-SpyVLP had at least one order of magnitude higher ACE2 blocking activity compared to serum from pigs, despite neutralisation titres being comparable (Figure 3D and 4D). This suggests that mice and pigs may produce distinct antibody responses against the vaccine candidate, although they were equally potent in neutralising live viruses. Surprisingly, ACE2 blocking in both mild and critical/severe convalescent humans who had natural infection were also low compared to the sera from immunised mice (Figure 3D). Nevertheless, the RBD-SpyVLP induces strong neutralising antibody responses in both mice and pigs. The potential for a vaccine based on the RBD is further emphasised by the sterile immunity induced in non-human primates (*Macaca mulatta*) by two doses of 20 to 40 µg of unconjugated RBD in Al(OH)_3_ adjuvant ^14^, and the successful induction of high titre neutralising antibody responses with an elegant self-assembling RBD-virus like nanoparticle ^50^.

A recently published report on an inactivated SARS-CoV-2 vaccine candidate showed that vaccinated non-human primates (NHP) with a serum neutralising titre (ND_50_) of <1:100 were still protected against wild type SARS-CoV-2 challenge with no weight loss and no detectable lung pathology ^51^. In our study, RBD-SpyVLP-vaccinated mice and pigs had ND_50_ at least an order of magnitude higher than 1:100, which would be expected to provide protection. Sera from pigs immunised with RBD-SpyVLP had similar, if not higher, neutralisation titres against wild type SARS-CoV-2 compared to pigs immunised by adenoviral vector (ChAdOx1 nCoV-19; both vaccines given as 2 doses in the same timeframe) ^34^. A single dose of ChAdOx1 nCoV-19 has been shown to be protective against viral pneumonia and lung pathology following SARS-CoV-2 challenge (ND_50_ ∼1:20) in NHP but not virus shedding in the nasal cavity ^52^. The seemingly similar ND_50_ of the RBD-SpyVLP compared to the ChAdOx1 nCoV-19 in pigs suggests that the RBD-SpyVLP will be as protective as the ChAdOx1 nCoV-19 in NHP and most likely in humans if similar neutralisation titres are achieved. A pre-stabilised spike glycoprotein vaccine candidate, NXV-CoV2273 developed by Novavax, showed neutralising antibody responses in humans that were 4-fold higher than convalescent human sera, when given two doses of 5 µg with Matrix M1 adjuvant ^53^. In light of the higher efficacy of RBD-SpyVLP compared to spike glycoprotein in pigs (Figure 4), we are hopeful that at least similar efficacy of RBD-SpyVLP could be achieved in humans.

Recently published studies reveal the relatively short-lived antibody response to SARS-CoV-2 in convalescent patients ^54, 55^. Further work is required to define the significance of these results for the longevity of protective immunity. It is possible that re-infection with the virus would lead to a strong protective secondary antibody response from memory B cells. Here, we have shown that the antibody response in pigs immunised with RBD-SpyVLP persisted for at least 2 months and remained neutralising. Studies on the longevity of the immune responses are to be undertaken in the near future.

Apart from being highly immunogenic, SpyVLPs provide a versatile modular vaccine platform to facilitate conjugation with other antigens. Should a mutation in the RBD arise in circulating SARS-CoV-2 strains ^48^, a matched RBD-SpyVLP could be manufactured rapidly. In addition, more than one RBD variant can be co-displayed on the VLP, to provide broader protection against various SARS-CoV-2 strains ^56^. The RBD-SpyVLP can also be co-displayed with antigens from other pathogens such as the HA and NA from influenza virus. We have recently shown HA and NA to be highly immunogenic in mice after formulation as a SpyVLP ^18^. This assembly could potentially provide protection against both SARS-Cov-2 and influenza viruses. Testing resilience of the vaccine candidate, we found that RBD-SpyVLP is stable at ambient temperature, resistant to freeze-thaw, and can be lyophilised and reconstituted with minimal loss in activity (Figure 2B-G) or immunogenicity (Figure 3F). This resilience may not only simplify vaccine distribution worldwide, especially to countries where cold-chain resources are incomplete, but also reduce the overall vaccine cost by removing cold-chain dependence. We are currently investigating cheaper and more scalable alternatives to produce RBD-SpyVLP to cope with the global demand for a SARS-CoV-2 vaccine. Collectively, our results show that the RBD-SpyVLP is a potent and adaptable vaccine candidate for SARS-CoV-2.

## MATERIALS AND METHODS

### Expression constructs

The SpyTag-RBD expression construct (Figure S1) consists of influenza H7 haemagglutinin (A/HongKong/125/2017) signal-peptide sequence, SpyTag ^57^, (GSG)_3_ spacer and SARS-CoV-2 spike glycoprotein (GenBank: NC045512) (amino acid 340-538). The insert was ordered from GeneArt and subcloned into pcDNA3.1 expression plasmid using unique *Not*I-*EcoR*I sites to create pcDNA3.1-SpyTag-RBD (GenBank and Addgene deposition in progress) (see Figure S1). pET28a-SpyCatcher003-mi3 (GenBank and Addgene deposition in progress) was created by replacing SpyCatcher in pET28a-SpyCatcher-mi3 with SpyCatcher003 ^18^. RBD used in the RBD ELISA was expressed from a codon optimised RBD cDNA subcloned into the vector pOPINTTGNeo incorporating a C-terminal His_6_ tag (RBD-6H) as previously described ^34^. Human ACE2 fused to human IgG1 Fc domain (ACE2-Fc) used in the ACE2 competition ELISA was expressed from codon optimised human ACE2 cDNA (amino acid 18 to 615) fused to the Fc region and a C-terminal His_6_ tag subcloned into the vector pOPINTTGNeo.

### Expression and purification of SpyCatcher003-mi3

SpyCatcher003-mi3 was expressed in *E. coli* BL21(DE3) RIPL cells (Agilent) as previously described ^18^. Heat-shock transformed cells were then plated on LB-Agar plates (50 µg/mL kanamycin) and incubated for 16 h at 37 °C. A single colony was picked and cultured in 10 mL starter LB culture (50 µg/mL kanamycin) for 16 h at 37 °C and shaking at 200 rpm. The preculture was diluted 1:100 into 1 L LB (50 µg/mL kanamycin and 0.8% (w/v) glucose) and cultured at 37 °C, 200 rpm until OD600 ∼0.6. Protein expression was induced with isopropyl β-D-1-thiogalactopyranoside (IPTG) (420 µM) and incubated at 22 °C, 200 rpm for a further 16 h. The culture was centrifuged and the pellet was resuspended in 20 mL 25 mM Tris-HCl, 300 mM NaCl, pH 8.5 with 0.1 mg/mL lysozyme, 1 mg/mL cOmplete mini EDTA-free protease inhibitor (Merck) and 1 mM phenylmethanesulfonyl fluoride (PMSF). Cell suspension was incubated at 22 °C for 30 min on a platform shaker and sonicated on ice 4 times for 60 s at 50% duty-cycle using an Ultrasonic Processor (Cole-Parmer). Cell lysate was clarified at 35,000 *g* for 45 min at 4 °C. The supernatant was filtered through 0.45 µm and 0.22 µm syringe filters (Starlab) and 170 mg ammonium sulfate was added per mL of lysate. SpyCatcher003-mi3 particles were precipitated by incubating the lysate at 4 °C for 1 h while mixing at 100 rpm. Precipitated particles were pelleted by centrifugation at 30,000 *g* for 30 min at 4 °C. The collected pellet was resuspended into 8 mL TBS pH 8.5 (25 mM Tris-HCl, 150 mM NaCl). Residual ammonium sulfate was removed by dialysing for 16 h against 500-fold excess of TBS. Dialysed SpyCatcher003-mi3 was concentrated to 4 mg/mL using a Vivaspin 20 100 kDa spin concentrator (Vivaproducts) and centrifuged at 17,000 *g* for 30 min at 4 °C to pellet any insoluble material. The supernatant was filtered through a 0.22 µm syringe filter. The purified SpyCatcher003-mi3 was then further purified using size exclusion chromatography (SEC). In brief, 2.5 mL was loaded into a HiPrep Sephacryl S-400 HR 16-600 SEC column (GE Healthcare) equilibrated with TBS using an ÄKTA Pure 25 system (GE Healthcare). Proteins were separated at 1 mL/min while collecting 1 mL elution factions. The fractions containing the purified particles were identified by SDS-PAGE, pooled, and concentrated using a Vivaspin 20 100 kDa MW cut-off centrifugal concentrator. Endotoxin was removed from the SpyCatcher003-mi3 samples using Triton X-114 phase separation as previously described ^18^. The concentration of endotoxin-depleted particles was measured using bicinchoninic acid (BCA) assay (Pierce) and particles were stored at −80 °C.

### Expression and purification of SpyTag-RBD

SpyTag-RBD was expressed in Expi293F cells using ExpiFectamine293 transfection reagent (Thermo Fisher) according to the manufacturer’s protocol. Supernatant was harvested between 5 to 7 days post transfection and filtered through a 0.22 µm filter, before purifying using Spy&Go affinity purification with minor modifications ^18, 23^. Briefly, filtered supernatants were diluted with one third supernatant volume of TP buffer (25 mM orthophosphoric acid adjusted to pH 7.0 at 22 °C with Tris base) and adjusted to pH 7. Spy&Go resin in same buffer was mixed with the diluted supernatant and incubated at 4 °C for 1 h with gentle agitation. The mixture was poured into an Econo-Pak column (Bio-Rad) and allowed to empty by gravity. The resin was washed with 2 × 10 column volumes of TP buffer and SpyTag-RBD was eluted with 2.5 M imidazole in TP buffer adjusted to pH 7.0 at room temperature (RT). One column volume of elution buffer was added to the resin at a time and incubated for 5 min before collecting each fraction. Elution fractions were analysed using SDS-PAGE with Coomassie staining and the fractions containing SpyTag-RBD were pooled and dialysed against 10 mM Tris-HCl pH 8.0 with 200 mM NaCl. The sample was concentrated using Vivaspin-20 10 kDa and further purified via SEC using ÄKTA Pure 25 (GE Life Sciences) equipped with Superdex 75pg 16-600 column (GE Life Sciences), run at 1 mL/min. The dialysis buffer was used as the mobile phase. The final yield of purified SpyTag-RBD was around 100 mg/L. The heterogeneity in the SpyTag-RBD band on SDS-PAGE is expected from the presence of different glycoforms on the N-linked glycosylation sites in the construct (Figure S1).

### RBD-SpyVLP conjugation

SpyTag-RBD at 2 µM, 3 µM, 4 µM or 6 µM were conjugated at 4 °C for 16 h with 2 µM SpyCatcher003-mi3 (VLP:RBD ratio 1:1, 1:1.5, 1:2 and 1:3) in TBS pH 8.0. Possible aggregates were then removed by centrifugation at 16,900 *g* for 30 min at 4 °C. Samples of the supernatant were mixed with reducing 6× loading dye (0.23 M Tris-HCl, pH 6.8, 24% (v/v) glycerol, 120 μM bromophenol blue, 0.23 M SDS, 0.2 M dithiothreitol) and resolved on 12%, 14% or 16% SDS-PAGE using XCell SureLock system (Thermo Fisher). Gels were then stained with InstantBlue Coomassie (Expedion) and imaged using ChemiDoc XRS imager (Bio-Rad). The intensities of bands on each lane were quantified using ImageLab (version 5.2) software (Bio-Rad) and Fiji distribution of ImageJ (version 1.51n). Conjugation efficiency (as % of unconjugated SpyVLP left) was calculated as (band density of unconjugated SpyVLP left in the conjugation reaction/band density of SpyVLP only control (2 μM)).

### RBD-SpyVLP thermostability and lyophilisation tests

30 μL of RBD-SpyVLP stored in thin-walled PCR tubes were subjected to freeze-thaw cycles (one to five cycles) by storing the tubes in a −80 °C freezer for 15 min or until the whole tube had frozen over, followed by incubation at RT for 10 min. For the storage temperature study, 30 μL of RBD-SpyVLP stored in thin-walled PCR tubes were incubated at −80 °C, −20 °C, 4 °C or RT (25 °C) for 14 days. The RBD-SpyVLP samples were then resolved on 4-12% Tris-Bis SDS-PAGE (Thermo Fisher) and analysed by densitometry following Quick Coomassie (Generon) staining. All samples were analysed in triplicate and plotted as mean ± 1 standard deviation (SD). The sample stored at −80 °C for two weeks, which had been through only 1 freeze-thaw cycle, was defined as 100% soluble. For lyophilisation, 100 μL of RBD-SpyVLP (125.5 μg/mL) in TBS pH 8.0 prepared in a Protein LoBind microcentrifuge tube (Fisher Scientific) was snap-frozen in liquid nitrogen for 30 min. A BenchTop 2K freeze-dryer (VisTis) was used for 24 h at 0.14 mbar and −72.5 °C to freeze-dry the sample. Lyophilised sample was reconstituted in the same original volume (100 μL) of MilliQ water and centrifuged at 16,900 *g* for 30 min to remove aggregates, before analysis with SDS-PAGE or ELISA. For testing of RBD-SpyVLP on ELISA, 50 μL of RBD-SpyVLP samples diluted in PBS (137 mM NaCl, 2.7 mM KCl, 10 mM Na2HPO4, 1.7 mM KH_2_PO_4_, pH 7.4) to 0.5 μg/mL were coated on NUNC plates at 4 °C overnight, washed with PBS, and blocked with 300 μL of 5% skimmed milk in PBS for 1 h at RT. Plates were then washed and incubated with 50 μL CR3022 (10 μg/mL) antibody (for freeze-thaw and storage temperature study) or a panel of mAbs as indicated in the graph (5 μg/mL) (for the lyophilisation study) diluted in PBS/0.1% BSA for 1 h at RT. Plates were washed and incubated with horse radish peroxidase (HRP) conjugated goat-anti-human IgG antibody (Dako, P0447) (diluted 1:1,600 in PBS/0/1% BSA) for 1 h at RT. Plates were then washed and developed with 50 μL of POD 3,3’,5,5’-tetramethylbenzidine (TMB) substrate (Roche) for 5 min and stopped with 50 μL of 1 M H_2_SO_4_. Absorbance was measured on a Clariostar plate reader (BMG Labtech). To test the reactivity of RBD-SpyVLP against a panel of anti-SARS-CoV-2 RBD antibodies, 50 μL of RBD-SpyVLP samples diluted in PBS to 0.5 μg/mL were coated on NUNC plates at 4 °C overnight. Plates were then washed and blocked with 300 μL of 5% (w/v) skimmed milk in PBS for 1 h at RT. Plates were washed and incubated with antibodies diluted in PBS with 0.1% (w/v) BSA in a 2-fold dilution series in 50 μL for 1 h. Second layer antibody was added as described above and plates were developed as above.

### Dynamic light scattering (DLS)

Samples were centrifuged for 30 min at 16,900 *g* at 4 °C to pellet possible aggregates. Before each measurement, the quartz cuvette was incubated in the instrument for 5 min to stabilise the sample temperature. Samples were measured at 125-250 µg/mL total protein concentration. 30 µL of sample was measured at 20 °C using an Omnisizer (Victotek) with 20 scans of 10 s each. The settings were 50% laser intensity, 15% maximum baseline drift, and 20% spike tolerance. The intensity of the size distribution was normalised to the peak value and plotted in GraphPad Prism 8 (GraphPad Software).

### Mouse immunisation and sampling

To prepare the RBD-SpyVLP for vaccination at 125 µg/mL (based on SpyTag-RBD concentration), 5 µM SpyTag-RBD was conjugated with 3.33 µM or 5 µM of SpyCatcher003-mi3 in TBS pH 8.0 at 4 °C for 16 h. The reaction was centrifuged for 30 min at 17,000 *g* at 4 °C to remove potential aggregates. RBD-SpyVLP was aliquoted and stored at −80 °C. Matching non-conjugated SpyTag-RBD or SpyCatcher003-mi3 VLP were diluted in the same buffers and incubated and centrifuged in the same way. RBD-SpyVLP (125 µg/mL) was diluted to 4 µg/mL (0.1 µg dose) or 20 µg/mL (0.5 µg dose) in the same buffer freshly before immunisation. Before IM immunisation, RBD-SpyVLP was mixed 1:1 (25 µL + 25 µL) with AddaVax adjuvant (Invivogen). Mouse experiments were performed according to the UK Animals (Scientific Procedures) Act Project Licence (PBA43A2E4) and approved by the University of Oxford Local Ethical Review Body. Female C57BL/6 or BALB/c mice (∼ 5 weeks old at the time of first immunisation) were obtained from BMS Oxford or Envigo. Mice were housed in accordance with the UK Home Office ethical and welfare guidelines and fed on standard chow and water ad libitum. Isoflurane (Abbott) lightly anaesthetised mice were immunised on day 0 and day 14 IM with 50 µL of RBD-SpyVLP at 0.1 µg or 0.5 µg or equivalent dose of unconjugated SpyTag-RBD or SpyCatcher003-mi3 VLP. Sera samples were obtained on day 42 via cardiac puncture of humanely sacrificed mice. The collected whole blood in Microtainer SST tubes (BD) was allowed to clot at RT for 1 h before spinning down at 10,000 *g* for 5 min. The clarified sera were heat-inactivated at 56 °C for 30 min before storing at −20 °C.

### RBD ELISA (Mouse and human sera)

RBD-6H was expressed in Expi293 according to the manufacturer’s protocol and purified using HisTrap HP column (Cytivia) and desalted using Zeba Spin Desalting Column (Thermo Fisher). To detect anti-RBD antibody in the immunised mouse sera, 50 µL purified RBD-6H (amino acids 330 to 532) (2 µg/mL) diluted in PBS was coated on NUNC plates at 4 °C overnight. Plates were then washed with PBS and blocked with 300 μL of 5% skimmed milk in PBS for 1 h at RT. In round-bottom 96-well plates, heat-inactivated mouse sera (starting dilution 1 in 40) was diluted in PBS/0.1% BSA in a 2-fold serial dilution in duplicate. 50 μL of the diluted sera was then transferred to the NUNC plates for 1 h at RT. Plates were then washed with PBS and 50 μL of secondary HRP goat anti-mouse antibody (Dako P0417) diluted 1:800 in PBS/0.1% BSA was added to the wells for 1 h at RT. Plates were washed and developed as described above. Serum RBD-specific antibody response was expressed as endpoint titre (EPT). EPT is defined as the reciprocal of the highest serum dilution that gives a positive signal (blank+10 SD) determined using a five-parameter logistic equation calculated using GraphPad Prism 8. RBD antibody response in the convalescent plasma was tested on a cell-based RBD ELISA, since human plasma gives high background on NUNC plates in our hands. MDCK-RBD cells were generated as previously described ^26, 39, 58^. Briefly, MDCK-SIAT1 cells (ECACC 05071502) ^59^were stably transfected using lentiviral vector to express SARS-CoV-2 RBD (amino acid 340-538, NITN…GPKK) fused at the C-terminus to the transmembrane domain of haemagglutinin H7 (A/HongKong/125/2017) (EPI977395) (KLSSGYKDVILWFSFGASCFILLAIVMGLVFICVKNGNMRCTICI) for surface expression. MDCK-RBD was seeded at 3 × 10^4^ cells/well in flat-bottom 96-well plates and incubated overnight at 37 °C and 5% CO_2_ prior to the assays. Human plasma was incubated with MDCK-SIAT1 cells for 1 h at RT prior to the assays to remove background binding on MDCK-SIAT1 cells. Plates seeded with MDCK-RBD were washed with PBS and 50 μL of pre-absorbed human sera diluted in a 2-fold dilution series (starting dilution 1 in 5) were added to the cells for 1 h at RT. The same set of sera were added in parallel plates seeded with MDCK-SIAT1 to obtain background binding on MDCK-SIAT1. Plates were washed and 50 μL of a secondary Alexa Fluor 647 goat anti-human antibody (Life Technologies A21455) (1:500 in PBS with 0.1% (w/v) BSA) were added for 1 h at RT. Plates were washed and 100 μL of PBS/1% formalin was added. Fluorescence signal was then read on a Clariostar plate reader. Background signal obtained on the parallel MDCK-SIAT1 plates were subtracted. EPT was determined as described above. Convalescent plasma samples were collected from March to May 2020 at John Radcliffe Hospital, Oxford. Patients’ severity was determined according to the WHO guidance (described in ^60^).

### Spike glycoprotein ELISA (Mouse and human sera)

A cell-based ELISA as described previously ^26^ was used to determine the anti-spike glycoprotein antibody response in the mouse sera and convalescent plasma. Briefly, MDCK-Spike was produced by stably transfecting parental MDCK-SIAT1 cells with full-length SARS-CoV-2 spike glycoprotein cDNA using a lentiviral vector. MDCK-Spike cells (3 × 10^4^ cells/well) was seeded in 96-well plates and incubated overnight at 37 °C. Mouse sera was diluted as above and 50 μL was transferred to the washed plates seeded with MDCK-Spike cells for 1 h at RT. Human plasma was pre-incubated with MDCK-SIAT1 cells as described above, before dilution and adding to MDCK-Spike for 1 h at RT. Parallel plates seeded with MDCK-SIAT1 for background subtraction was done as described above for human plasma. For mouse sera, 50 μL of a secondary Alexa Fluor 647 goat-anti mouse antibody (1:500) (Life Technologies A21235) was then added for 1 h at RT. For human sera, 50 μL of a secondary Alexa Fluor 647 goat-anti human antibody (1:500) (Life Technologies A21455) was used. Plates were then washed with PBS and 100 μL of PBS/1% formalin was added to each well. Fluorescence signal was read on a Clariostar plate reader and the EPT titre was calculated as described above.

### ACE2 competition ELISA

ACE2 competition ELISA was done with RBD-SpyVLP immobilised on NUNC plates as previously described with slight modifications ^26^. ACE2-Fc was expressed in Expi293 and purified using MabSelect SuRe column (Cytivia) and salts were removed using Zeba Spin Desalting Column (Thermo Fisher) into PBS. ACE2-Fc was chemically biotinylated using EZ-Link Sulfo-NHS-LC-Biotinylation Kit (Thermo Fisher) according to manufacturer’s protocol. 25 ng/well of RBD-SpyVLP was coated on the ELISA plate at 4 °C overnight. Plates were then washed with PBS and blocked with 300 μL of 5% (w/v) skimmed milk in PBS for 1 h at RT. Heat-inactivated mouse or pig sera or human plasma samples were titrated in duplicate as half-log_10_ 8-point serial dilution, starting at 1 in 5 in 30 µL with PBS with 0.1% (w/v) BSA. 30 µL of biotinylated ACE2-Fc at 0.2 nM (40 ng/mL) (or 0.4 nM for human plasma) was added to the samples. 50 µL of the biotinylated ACE2-Fc:sample mixture was transferred to the RBD-SpyVLP coated plates and incubated for 1 h at RT. A secondary Streptavidin-HRP (S911, Life Technologies) diluted to 1:1,600 in PBS/0.1% BSA was added to the PBS-washed plates and incubated for 1 h at RT. Plates were then washed and developed as above. Biotinylated ACE2-Fc without sera or plasma was used to obtain the maximum signal and wells with PBS/BSA buffer only were used to determine the minimum signal. Graphs were plotted as % binding of biotinylated ACE2 to RBD. Binding % = (X - Min) / (Max - Min) * 100 where X = measurement of the sera or plasma, Min = buffer only, Max = biotinylated ACE2-Fc alone. ACE2 blocking activity of the sera or plasma was expressed as IC_50_ determined using non-linear regression curve fit using GraphPad Prism 8.

### Authentic SARS-CoV-2 virus neutralisation assay (PRNT)

96 well plates containing a confluent monolayer of Vero-E6 cells were incubated with 10-20 plaque forming units (PFU) of SARS CoV-2 (hCoV-19/England/02/2020, EPI_ISL_407073, kindly provided by Public Health England) and two-fold serial dilution of heat-inactivated mouse sera for 3 h at 37 °C, 5% CO_2_, in triplicate per serum sample. Inoculum was then removed, and cells were overlaid with virus growth medium containing Avicel microcrystalline cellulose (final concentration of 1.2%) (Sigma-Aldrich). The plates were then incubated at 37 °C, 5% CO_2_. At 24 h post-infection, cells were fixed with 4% paraformaldehyde and permeabilised with 0.2% (v/v) Triton-X-100 in PBS stained to visualise virus plaques, as described previously for the neutralisation of influenza viruses ^61^, but using a rabbit polyclonal anti-NSP8 antibody (Antibodies Online; ABIN233792) and anti-rabbit-HRP conjugate (Bio-Rad) and detected using HRP on a TMB based substrate. Virus plaques were quantified and ND_50_ for sera was calculated using LabView software as described previously ^61^.

### Authentic SARS-CoV-2 plaque reduction neutralization assay (PRNT)

SARS-CoV-2 (hCoV-19-Australia/VIC01/2020, GenBank MT007544) ^62^ was diluted to a concentration of around 1,000 PFU/mL and 75 μL was mixed with an equal volume of minimal essential medium (MEM) (Life Technologies) containing 1% (v/v) foetal bovine serum (FBS) (Life Technologies) and 25 mM HEPES buffer (Sigma-Aldrich) with doubling pooled mouse sera dilutions (starting dilution 1:40) in a 96-well V bottomed plate. The plate was then incubated at 37 °C in a humidified incubator for 1 h before the virus-antibody mixture was transferred 24-well plates containing confluent monolayers of Vero E6 cells (ECACC 85020206) cultured in MEM containing 10% (v/v) FBS. The plates were incubated for 1 h at 37 °C and overlaid with MEM containing 1.5% carboxymethylcellulose (Sigma-Aldrich), 4% (v/v) FBS and 25mM HEPES buffer. Plates were incubated at 37 °C for five days prior to fixation with 20% formalin in PBS overnight. Plates were then washed with tap water and stained with 0.2% crystal violet solution (Sigma-Aldrich) and plaques were visualised and counted. A mid-point probit analysis (written in R programming language for statistical computing and graphics) was used to determine the dilution of antibody required to reduce numbers of SARS-CoV-2 virus plaques by 50% (ND_50_) compared with the virus-only control (n = 5). A human MERS convalescent serum known to neutralise SARS-CoV-2 (National Institute for Biological Standards and Control, UK) was included in each run as assay control.

### mAb competition assays

mAb competition ELISA was done with RBD-SpyVLP immobilised on NUNC plates as described above with slight modifications. Plates were coated and blocked as above. Heat-inactivated mouse or pig sera or human plasma samples were titrated in duplicate as half-log 8-point serial dilution, starting at 1 in 5 in 30 µL with PBS with 0.1% (w/v) BSA or tested at 1:20 in quadruplicates. 30 µL of chemically biotinylated mAb (FI-3A, FD-11A, EY6A or S309) titrated to determine the lowest binding concentration at top plateau (all mAbs produced in house) ^26, 38, 39^ was added to the sera. Biotinylation was conducted as described above. 50 µL of the biotinylated mAb:sera mixture was transferred to the RBD-SpyVLP coated plates and incubated for 1 h at RT. A secondary Streptavidin-HRP (Life Technologies, S911) diluted to 1:1,600 in PBS/0.1% BSA was added to the PBS-washed plates and incubated for 1 h at RT. Plates were then washed and developed as above. Biotinylated mAb without sera or plasma was used to obtain the maximum signal and wells with PBS with 0.1% (w/v) BSA buffer only were used to determine the minimum signal. Graphs were plotted as % binding of biotinylated mAb to RBD-SpyVLP. Binding % = (X - Min) / (Max - Min) * 100 where X = measurement of the sera or plasma, Min = buffer only, Max = biotinylated mAb alone.

### Pig immunisation and sampling

Pig studies were performed in accordance with the UK Animals (Scientific Procedures) Act 1986 and with approval from the local Animal Welfare and Ethical Review Body (AWERB) (Project Licence PP1804248). RBD-SpyVLP for pig immunisation was prepared as above. Nine weaned, Large White-Landrace-Hampshire cross-bred pigs of 8–10 weeks of age from a commercial rearing unit were randomly allocated to three treatment groups (5 µg RBD-SpyVLP, 50 µg RBD-SpyVLP or 100 µg spike glycoprotein) (n = 3). RBD-SpyVLP was diluted to 5 µg/mL or 50 µg/mL in 25 mM Tris-HCl, pH 8.0, 150 mM NaCl and mixed with an equal amount of AddaVax (Invivogen) (1 mL + 1 mL) prior to immunisation. The spike glycoprotein was a soluble trimeric spike with the pre-fusion stabilisation substitutions (K983P, V984P, furin cleavage site removed and inclusion of a C-terminal T4-foldon domain for trimerization) ^63^. The spike glycoprotein was expressed in Expi293 cells and purified as previously described ^34^. Briefly, supernatant containing soluble spike glycoprotein was purified using immobilised metal affinity followed by gel filtration in TBS (pH 7.4). Pigs were dosed via IM injection into the brachiocephalic muscle with 2 mL of RBD-SpyVLP (5 µg or 50 µg) or spike (100 µg) at day 0 and day 28. Blood samples were taken on a weekly basis at 0, 7, 14, 21, 28, 35, 42 and 56 days post-immunisation (DPI) by venepuncture of the external jugular vein: 8 mL/pig in BD SST vacutainer tubes (Fisher Scientific) for serum collection and 40 mL/pig in BD heparin vacutainer tubes (Fisher Scientific) for PBMC isolation. Additional heparin blood samples were collected on 31 and 33 DPI to track the plasma cell response to boost. Sera samples were stored at −20 °C and heat-inactivated at 56 °C for 2 h before use in pVNT or VNT assays. Oral and nasal swabs were collected weekly and placed in 500 µL Media 199 (Thermo Fisher) supplemented with 0.0025% Nystatin (Merck), 0.01% Penicillin-Streptomycin (Gibco), 0.025% 1M HEPES solution (Gibco), 0.005% (w/v) sodium bicarbonate (Merck) and 0.067% (w/v) BSA (Merck) (VTM). Swabs were centrifuged at 700 × *g* for 5 min before aspirating the liquid and storing with the swab at −20 °C. Prior to assessment of antibodies, swabs and VTM were loaded in Spin-X Centrifuge 0.45 µM columns (Fisher Scientific) and fluid collected by centrifugation at 21,000 × *g* for 5 min.

### RBD ELISA (Pig sera and swab fluids)

An ELISA to analyse anti-RBD antibody response in pig sera was performed as previously described ^34^. Briefly, 50 µL of 2 µg/mL purified RBD-6H as described above was coated on flat-bottomed 96-well plates (Immunon 4 HBX; Thermo Fisher Scientific) overnight at 4 °C. Plates were washed with TBS (pH 7.4) with 0.1% (v/v) Tween-20 and blocked with 100 µL of PBS containing 3% skimmed milk for 1 h at RT. Pig sera samples were diluted in PBS with 1% (w/v) skimmed milk and 0.1% (v/v) Tween-20 in a 2-fold serial dilution starting at 1:10 dilution and 100 µL of the diluted sera was added to the coated plates for 1 h at RT. A conjugated secondary goat anti-pig IgG HRP (Abcam, Cambridge, UK) at 1:10,000 dilution in PBS with 1% (w/v) skimmed milk and 0.1% (v/v) Tween-20 was added for 1 h at RT. Plates were washed and 100 µL TMB (One Component Horse Radish Peroxidase Microwell Substrate, BioFX, Cambridge Bioscience) was added to each well and the plates were incubated for 7 min at RT. 100 µL BioFX 450nmStop Reagent (Cambridge Bioscience) was then added and absorbance was determined using a microplate reader. End-point antibody titres (mean of duplicates) were defined as following: the log_10_ OD was plotted against the log_10_ sample dilution and a regression analysis of the linear part of this curve allowed calculation of the endpoint titre with an absorbance of twice the mean absorbance of pre-immunised sera. RBD-specific antibody titres in oral and nasal swab fluids were determined by ELISA as detailed above except that the conjugated secondary antibody was replaced with either goat anti-porcine IgG HRP (Bio-Rad Antibodies) at 1:20,000 dilution in PBS with 1% (w/v) skimmed milk and 0.1% (v/v) Tween-20 or goat anti-porcine IgA HRP (Bio-Rad Antibodies) at 1:20,000 dilution in the same diluent.

### Virus neutralization assay (VNT) (pig sera)

VNT on pig sera was done as described previously ^34^. Briefly, Vero E6 cells were seeded in 96-well flat-bottom plates (1 × 10^5^ cells/mL) and incubated at 37 °C overnight prior to the assays. Two-fold serial dilutions of sera (starting dilution 1 in 5) in quadruplet were prepared in 96-well round-bottom plates using Dulbecco’s Modified Eagle Medium (DMEM) with 1% (v/v) FBS and 1% Antibiotic-Antimycotic (Gibco). 75 µL of the diluted sera was mixed with an equal volume of media containing 64 PFU of SARS-CoV-2 virus (hCoV-19/England/02/2020, EPI_ISL407073) and incubated for 1 h at 37 °C. Media in the wells seeded with Vero E6 was replaced with 100 µL DMEM with 10% (v/v) FBS and 1% Antibiotic-Antimycotic (Gibco) and 100 µL of the sera-virus mixture was added into the wells. The plates were incubated for six days at 37 °C. Cytopathic effect (CPE) was monitored on a brightfield microscopy, and by fixation using formaldehyde (VWR, Leighton Buzzard, UK) and staining using 0.1% Toluidine Blue (Sigma-Aldrich). CPE was scored by researchers who were blinded to the identity of the samples. No sera or no virus controls were run in parallel on each plate. Neutralisation titres (ND5_0_) were expressed as the reciprocal of the serum dilution that prevented CPE in 50% of the wells.

### Pseudovirus neutralisation test (pVNT) (pig sera)

Lentiviral-based SARS-CoV-2 pseudoviruses were generated as described previously ^34^. Briefly, HEK293T cells were seeded at a density of 7.5 × 10^5^ in 6-well plates before transfection with the following plasmids: 500 ng of SARS-CoV-2 spike, 600 ng p8.91 (HIV-1 gag-pol), 600 ng CSFLW (lentiviral genome plasmid encoding a firefly luciferase transgene) ^34^ using 10 µL polyethylene imine (PEI) (1 µg/mL) In Opti-MEM media (Thermo Fisher). The media was replaced with 3 mL DMEM with 10% FBS and incubated at 37 °C. Supernatant containing pseudovirus was harvested at 48 h and 72 h post transfection. Collected supernatant was centrifuged at 1,300 x *g* for 10 min at 4 °C to remove debris. To perform the assay, 2 × 10^4^ HEK293T target cells transfected with 500 ng of a human ACE2 expression plasmid (Addgene) were seeded in a white flat-bottom 96-well plate one day prior to the assays. Pig sera were diluted with a four-fold serial dilution in serum-free media (starting dilution 1 in 20) and 50 µL was added to a 96-well plate in quadruplicate. Pseudovirus pre-titrated to give 1 × 10^6^ relative light unit (RLU) in 50 µL DMEM with 10% FBS was added to the sera and incubated at 37 °C for 1 h. The pseudovirus:sera mix was then transferred to the target cells and incubated at 37 °C for 72 h. Firefly luciferase activity was measured using BrightGlo luciferase reagent on a GloMax-Multi+ Detection System (Promega). Pseudovirus neutralisation titres (ND_50_) were expressed as the reciprocal of the serum dilution that inhibited luciferase signal in 50% of the wells.

### Intracellular cytokine staining assay (pig PBMC)

Assessment of intracellular cytokine expression following stimulation of PBMCs with synthetic peptides representing SARS-CoV-2 S protein was conducted as described previously ^34^. In brief, PBMCs were isolated from heparinized blood by density gradient centrifugation and suspended at 1 × 10^7^ cells/mL in RPMI-1640 medium, GlutaMAX supplement, HEPES (Gibco) supplemented with 10% (v/v) heat-inactivated FBS (New Zealand origin, Life Science Production), 1% Penicillin-Streptomycin and 0.1% 2-mercaptoethanol (50 mM; Gibco) (cRPMI). 50 µL PBMC were added per well to 96-well round bottom plates and stimulated in triplicate wells with SARS-CoV-2 S peptide pools at a final concentration of 1 µg/mL peptide. Unstimulated cells in triplicate wells were used as a negative control. After 14 h incubation at 37°C, 5% CO_2_, cytokine secretion was blocked by addition 1:1,000 BD GolgiPlug (BD Biosciences) and cells were further incubated for 6 h. PBMC were surface-labelled with Zombie NIR fixable viability stain (BioLegend), CD3-FITC mAb (clone BB23-8E6-8C8, BD Biosciences), CD4-PerCP-Cy5.5 mAb (clone 74-12-4, BD Bioscience) and CD8α-PE mAb (clone 76-2-11, BD Bioscience). After fixation (Fixation Buffer, BioLegend) and permeabilization (Permeabilization Wash Buffer, BioLegend), cells were stained with: IFN-γ-Alexa Fluor 647 mAb (clone CC302, Bio-Rad Antibodies,) and TNF-α-Brilliant Violet 421 mAb (clone Mab11, BioLegend). Cells were analysed using a BD LSRFortessa flow cytometer (BD Biosciences) and data analysed using FlowJo software (BD Biosciences). Total SARS-CoV-2 S-specific IFN-γ-positive responses for live CD3^+^CD4^+^ and CD3^+^CD4^-^CD8^+^ T cells are presented after subtraction of the background response detected in the media-stimulated control PBMC samples of each pig, prior to summing together the frequency of S-peptide pools 1-3 specific cells.

### RBD-tetramer staining assay (pig PBMC)

A biotinylated form of RBD was generated for B-cell tetramer staining assays. An RBD protein with a C-terminal biotin acceptor peptide (RBD-BAP) was expressed from plasmid pOPINTTGNeo in Expi293 cells according to the manufacturer’s instructions. Culture supernatants were clarified by centrifugation and purified through a 5 mL HisTrap FF column (GE Healthcare), using the<u>Ä</u>KTA Pure chromatography system (Cytiva). Fractions containing RBD-BAP were concentrated and the excess imidazole removed by buffer exchange using an Amicon 10kDa (Merck). RBD-BAP was biotinylated using *E. coli* biotin ligase (BirA). GST-BirA enzyme was expressed, purified and biotinylated as previously described ^64^. Biotinylation reactions were assembled with 100µM RBD-BAP, 1µM GST-BirA, 5mM magnesium chloride (Ambion), 2mM ATP and 150µM D-Biotin (both Merck) and incubated twice for one hour at 30 °C with additional fresh biotin and GST-BirA added in-between incubations. GST-BirA was removed from the reaction with a GST HiTrap column, as above, and RBD-BAP was purified out by dialysis as above. Biotinylation of RBD-BAP was confirmed by streptavidin band shift assay ^64^ and quantified by BCA assay (Pierce). RBD tetramers were assembled by combining biotinylated RBD with streptavidin-Brilliant Violet 421 or streptavidin-Brilliant Violet 650 (both BioLegend) at a molar ratio of 4:1. Negative control ‘decoy’ tetramers were similarly assembled using biotinylated Nipah virus soluble glycoprotein ^65^ and streptavidin-PerCP Cy5.5 (BioLegend).

PBMC were stained with SARS-CoV-2 S RBD-tetramers to assess the frequency of circulating specific B cells during the course of the study. For 28, 31, 33 and 35 days post-infection, fresh PBMC were analysed (in triplicate), while for 0, 7, 14, 42 and 56 days post-infection, previously cryopreserved PBMC were assessed (in quadruplicate). RBD tetramers were assembled by combining biotinylated RBD with streptavidin-Brilliant Violet 421 or streptavidin-Brilliant Violet 650 (both BioLegend) at a molar ratio of 4:1. Negative control ‘decoy’ tetramers were similarly assembled using biotinylated Nipah virus soluble glycoprotein ^65^ and streptavidin-PerCP Cy5.5 (BioLegend). PBMC were washed in cold PBS and seeded at 1 × 10^6^ cells/well in 96-well round bottom plates and with combinations of RBD and decoy tetramers by incubation for 30 min on ice. After washing, cells were stained with Zombie Aqua, CD3-PE-Cy7 mAb (clone BB23-8E6-8C8, BD Biosciences), CD14-PE Vio 770 mAb (clone REA599, Miltenyi Biotec), IgG-Alexa Fluor-647 mAb (Cohesion Biosciences, Generon,), IgA-FITC polyclonal Ab (BioRad Antibodies) and IgM-PE mAb (clone K52 1C3, BioRad Antibodies; conjugated using Lightning-Link® PE Antibody Labeling Kit, Expedeon, Abcam), for 30 min on ice. After washing and fixation in 4% paraformaldehyde for 30 min at 4°C, cells were analysed on a BD LSRFortessa flow cytometer with downstream analysis using FlowJo software. Following exclusion of dead cells, CD3^+^, CD14^+^ and decoy tetramer+ cells, the percentage of IgA^+^, IgG^+^ or IgM^+^ cells dual-labelled with both RBD tetramers was assessed.

### RBD IgG ELISpot assay (pig PBMC)

Sterile 96-well Multiscreen-HA filter plates with a mixed cellulose membrane (MAHAS4510, Millipore) were coated with 100 μL of 15 μg/mL anti-porcine IgG mAb (clone MT421, Mabtech, 2BScientific) diluted in 0.05M carbonate-bicarbonate buffer pH 9.2 (Merck). Coated plates were incubated for a minimum of 18 h at 4 °C. Plates were then washed with PBS and blocked using cRPMI for at least 1 h at 37 °C, 5% CO_2_, 95% humidity. Blocking solution was then removed, and PBMC were added at a density of 5 × 10^5^/well for antigen-specific response or at 5 × 10^4^/well for wells assigned to total IgG (positive control). The plates were then incubated for 18 h at 37 °C, 5% CO_2_, 95% humidity. Media was removed and cells were lysed with cold distilled water, followed by three PBS washes as before. To measure total IgG, 50 μL/well of biotinylated anti-IgG mAb (clone MT424-biotin, Mabtech) was added at 0.5 µg/mL. To assess antigen-specific responses 50 μL/well of biotinylated SARS-CoV-2 RBD was added at 2.5μg/mL. As a negative control, 50 μL/well of biotinylated Nipah G protein was added to the relevant wells at 2.5 μg/mL. All antigens were diluted in PBS with 0.5% (v/v) FCS. An additional set of negative control wells were also prepared by adding 50 μL/well PBS with 0.5% (v/v) FCS. Each condition was tested in triplicate. Plates were incubated for 2 h at RT, before washing five times with PBS.

Following this, 50 μL/well of streptavidin-alkaline phosphatase (streptavidin-ALP) enzyme conjugate (Mabtech) (diluted 1:1,000 in PBS with 0.5% (v/v) FCS) was added to each well and plates were incubated for 1 h at RT (protected from light). Streptavidin-ALP was removed, and plates were washed another 5 times with PBS, followed by addition of 50 μL/well BCIP/NBTplus substrate (Mabtech), neat. Plates were left for 30 min, until distinct spots developed. Finally, development was stopped by addition of 150 μL/well of 4 °C distilled water followed by rinsing both the front and back of the plates with copious tap water. Plates were air-dried, before spots were counted using a CTL ImmunoSpot Analyzer (Cellular Technologies).

## Statistical analysis

All statistical analyses were performed using GraphPad Prism 8 (GraphPad Software). Statistical differences were analysed using either Mann-Whitney U test or Kruskal-Wallis test followed by Dunn’s multiple comparisons. A p value <0.05 was deemed statistically significant.

## CONTRIBUTIONS

**Production & characterisation of RBD-SpyVLP and mouse immunogenicity studies:** T.K.T. performed the RBD-SpyVLP conjugation, *in vitro* antigenicity and stability characterisation, and all mouse experiments and analysis. P.R. produced the RBD protein, developed mice and human immunoassays and performed analysis on mouse and pig sera. L.S. performed competition ELISA on mouse and pig sera. R.R. and A.H.K. produced and purified the SpyCatcher003 VLP, purified the RBD protein, and performed DLS and lyophilisation. T.K.T., P.R., M.H. and A.R.T. conceived and designed the experiments. KY.A.H provided the monoclonal antibodies. S.H., R.H., R.S.D. and J.W.M. designed and performed neutralisation assays on the mouse sera. J.A.T., K.R.B. and M.W.C performed the VNT on the pooled mouse sera. **Pig immunogenicity studies:** J.W.P.H., J.C.E., R.K.M, M.P., V.M., K.M., C.C., R.W., A.G., M.A., V.M., and S.P.G conducted the pig immunogenicity study, processed samples and conducted the T cell and B cell analyses. N.Z., C.C., M.T., and D.B. conducted the pVNT assays with pig sera. I.D., H.S., A.Z., D.B., S.B., P.S.L. and P.H. conducted the VNT assays with pig sera. A.L., G.W. and C.B. conducted the ELISAs with pig sera and swab fluids. J.N., A.S.A, A.B., S. C., T.M., J.H. and R.A. produced recombinant RBD, RBD-biotin and spike protein. R.W., J.A.H., E.T. B.C., T.J.T., and S.P.G. conceived and designed experiments. T.K.T. prepared the manuscript. All authors read, reviewed and approved the manuscript.

## ACKNOWLEDGEMENT AND FUNDING

T.K.T. is funded by the Townsend-Jeantet Charitable Trust (charity number 1011770) and the EPA Cephalosporin Early Career Researcher Fund. P.R., L.S. and A.R.T. are funded by the Chinese Academy of Medical Sciences (CAMS) Innovation Fund for Medical Science (CIFMS), China (grant no. 2018-I2M-2-002). The work done at the Crick Worldwide Influenza Centre was supported by the Francis Crick Institute, receiving core funding from Cancer Research UK (FC001030), the Medical Research Council (FC001030) and the Wellcome Trust (FC001030). The pig study was supported by UKRI Biotechnology and Biological Sciences Research Council (BBSRC) Institute Strategic Programme and Core Capability Grants to The Pirbright Institute (BBS/E/I/00007031, BBS/E/I/00007034, BBS/E/I/00007037 and BBS/E/I/00007039), and the Bill and Melinda Gates Foundation supported ‘Pirbright Livestock Antibody Hub’ (Grant No. OPP1215550). Development of SARS-CoV-2 reagents was partially supported by EPSRC Grant No. EP/S025243/1 to the Rosalind Franklin Institute. A.L., G.W., C.B., A.B., and V.M. are supported by the UK Department for Environment Food and Rural Affairs (Grant No. SE26081). We thank The Pirbright Institute Animal Services Team for animal care and provision of samples.

