## Supplementary for "A COVID-19 vaccine candidate using SpyCatcher multimerization of the SARS-CoV-2 spike protein receptor-binding domain induces potent neutralising antibody responses"

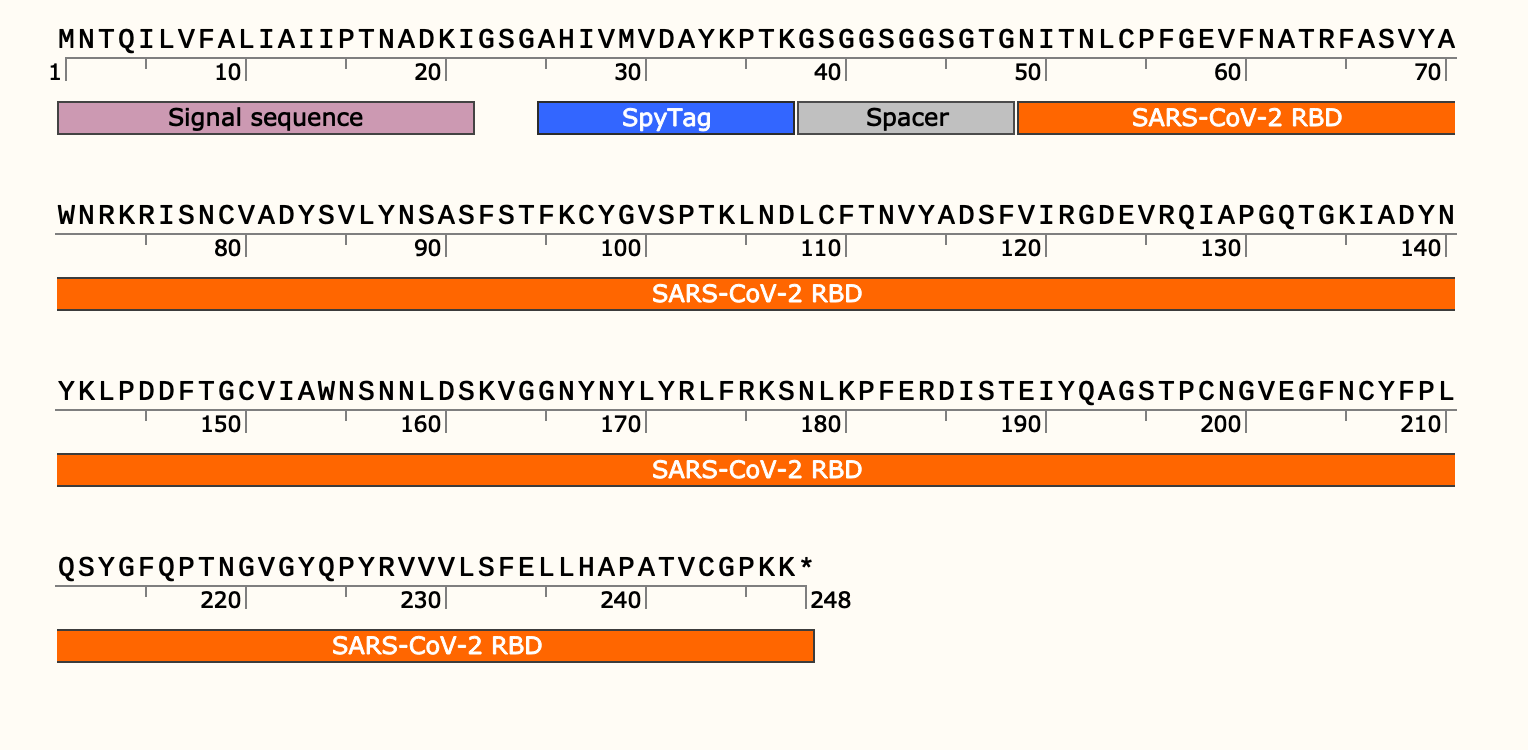

**Figure S1: SpyTag-RBD expression construct.** The expression cassette of SpyTag-RBD comprises the signal sequence from influenza H7 haemagglutinin (A/HongKong/125/2017) (mauve), SpyTag (blue), spacer (grey) and SARS-CoV-2 RBD 340-538 (orange). Diagram adopted from SnapGene.

**Figure S2: Analysis of the thermostability of RBD-SpyVLPs**. (A) RBD-SpyVLPs (n=3 for each temperature) were stored at the indicated temperatures for two weeks, prior to analysis on SDS-PAGE with Coomassie staining. Solubility was calculated using densitometry and presented in Figure 2B. (B) RBD-SpyVLPs (n=15, n=3 per condition) were subjected to 1-5 rounds of free-thaw cycles as indicated and analysed on SDS-PAGE with Coomassie staining. Solubility was calculated using densitometry and presented in Figure 2D. * trace of proteolysis product.

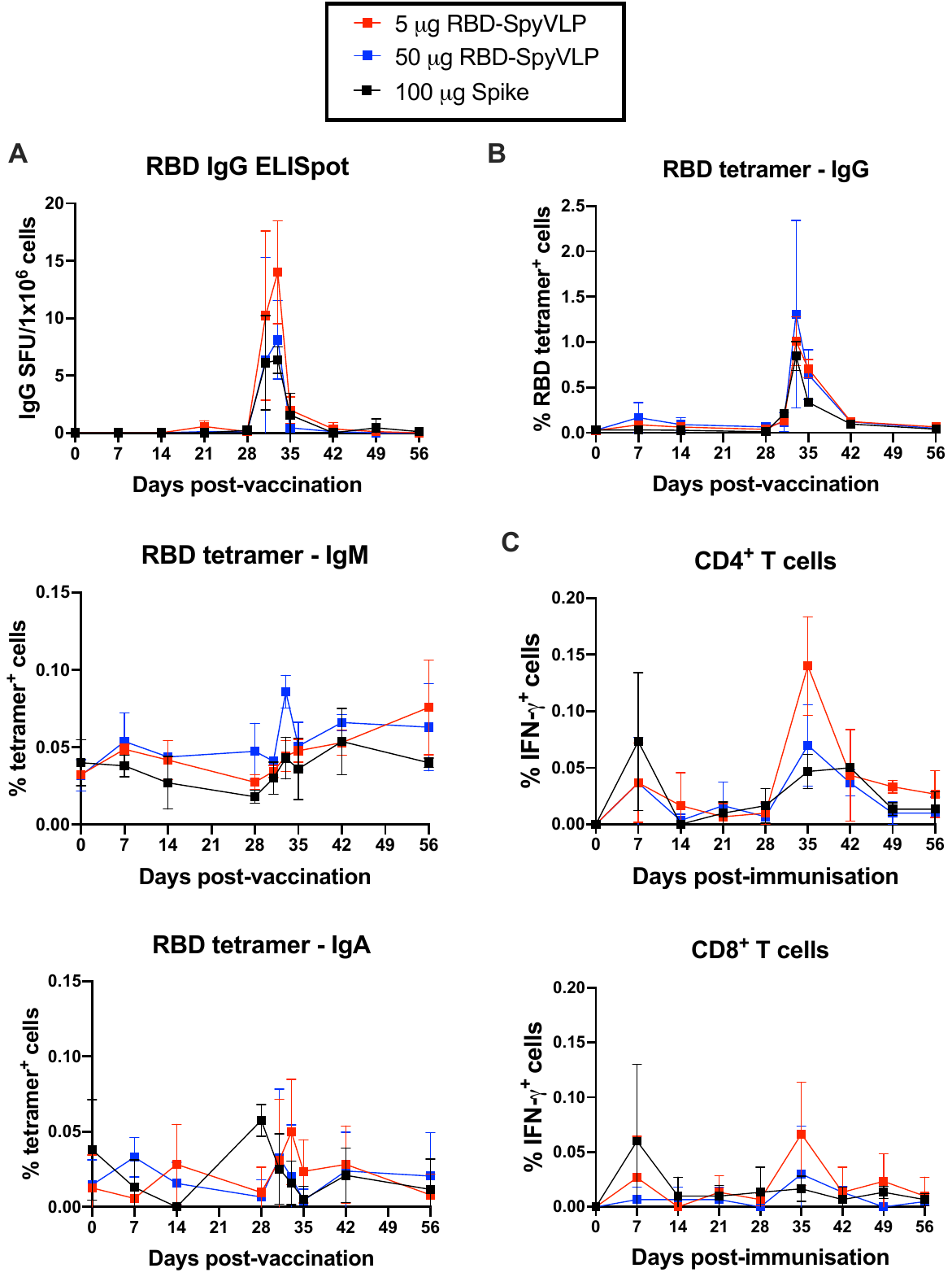

**Figure S3: B cell and T cell analyses of pigs immunised with RBD-SpyVLPs or Spike glycoprotein.**  SARS-CoV-2 S-specific B cell and T cell responses were longitudinally assessed in PBMC following immunisation of pigs with RBD-SpyVLP (5 µg or 50 µg) or 100 µg Spike glycoprotein on 0- and 28-days post-immunisation. (A) B cell responses were assessed by flow cytometric analysis of IgG^+^, IgM^+^ and IgA^+^ cells binding to RBD-tetramers. (B) RBD-specific plasma cell responses were assessed by IgG ELISpot assay. SFU = spot-forming units. (C) T cell responses were assessed by flow cytometry. Following stimulation with S peptides, IFN-γ expression by CD4^+^ and CD8^+^ T cells was determined by intracellular cytokine staining. Data points represent the mean ± 1 SEM (n=3).

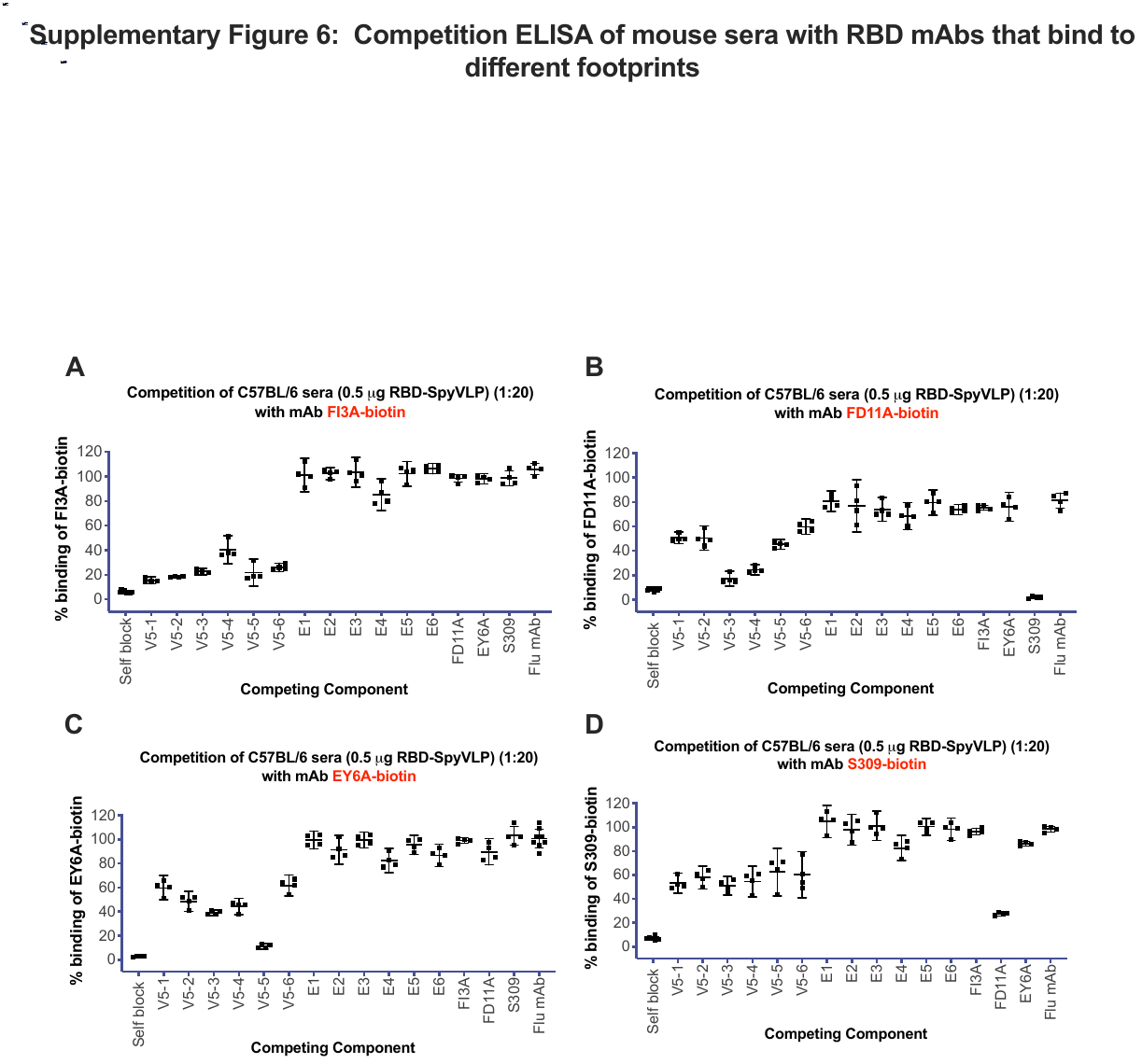

**Figure S4: Competition ELISA of C57BL/6 sera with RBD mAbs binding to non-overlapping epitopes on RBD**. Competition ELISA of C57BL/6 mice immunised with 0.5 µg RBD-SpyVLP (sera from 3 weeks post boost) against (A) FI-3A biotin, (B) FD11A biotin, (C) EY6A biotin and (D) S309 biotin. 1:20 serum dilutions were used and each serum was tested in quadruplicate. Data are shown as the mean of the readings ± 95% confidence interval.

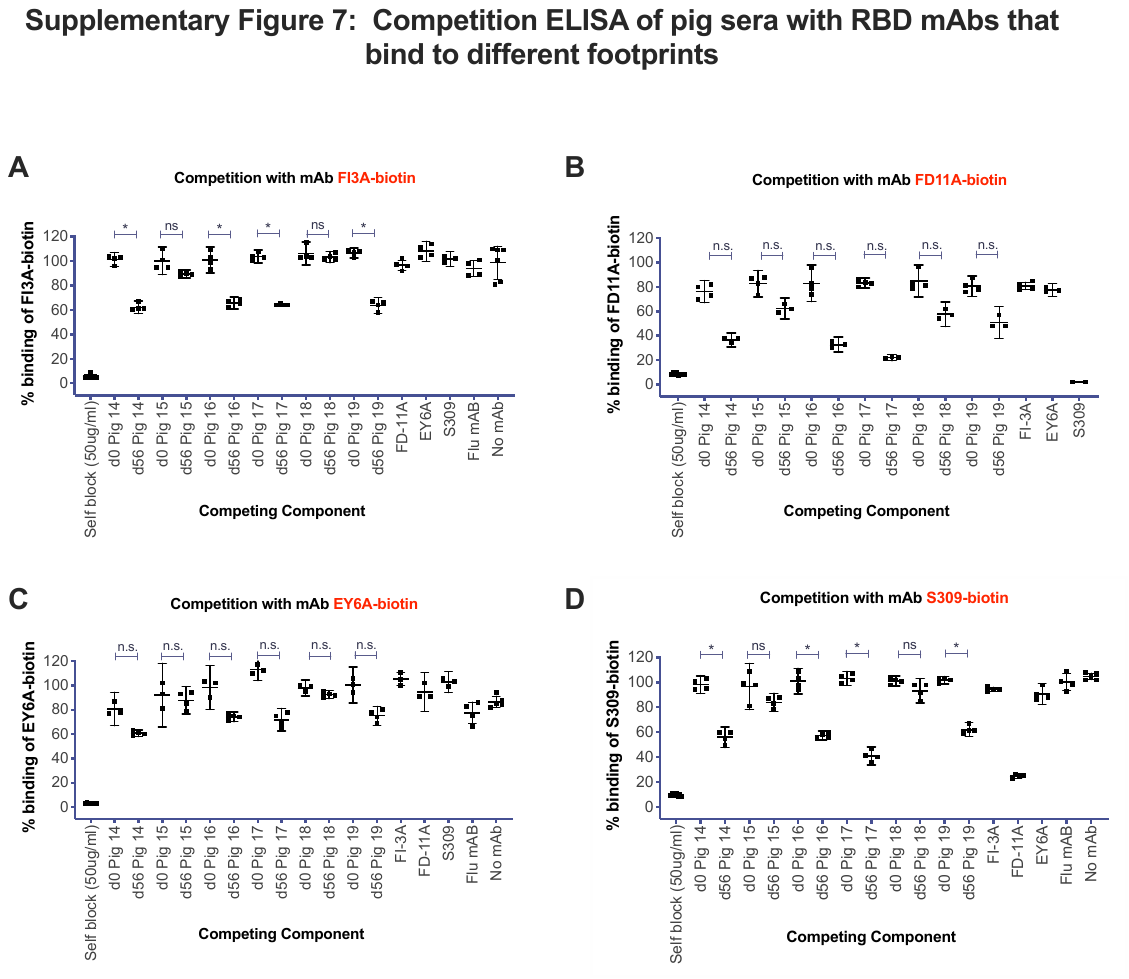

**Figure S5: Competition ELISA of pig sera with RBD mAbs binding to non-overlapping epitopes on RBD**. (A) Competition ELISA of preimmune sera (day 0) and post boost sera (day 56) for pigs (n=3) immunised with 5 µg RBD-SpyVLP (pig 14, 15 & 16) or 50 µg RBD-SpyVLP (pig 17, 18 & 19) against (A) FI-3A biotin, (B) FD11A biotin, (C) EY6A biotin and (D) S309 biotin. Data are shown as the mean of the readings ± 95% confidence interval. *p<0.05, Mann Whitney U test between day 56 and preimmune (day 0) sera samples. n.s = not significant

| **VNT on pooled sera (C57BL/6)** | | |
| --- | --- | --- |
| **Group** | **Probit mid-point PRNT_50_ dilution (ND_50_)** | **95% confident intervals** |
| **0.1 μg RBD** | 395 | 113-751 |
| **0.5 μg RBD** | 79 | 16-159 |
| **0.1 μg RBD-SpyVLP** | 6,087 | 4,487-9,375 |
| **0.5 μg RBD-SpyVLP** | 8,789 | 6,710-12,966 |
| **VLP** | 122 | 89-147 |
| **MERS control sera** | 527 | 419-651 |

**Table S1: VNT of pooled sera from C57BL/6 mice immunised with RBD-SpyVLP**. The table summarises the ND_50_ of pooled sera of C57BL/6 mice (n=6) immunised with RBD, RBD-SpyVLP or VLP against live SARS-CoV-2 in a plaque reduction-based neutralisation assay. A serum panel from humans infected with MERS-CoV, known to cross-neutralise SARS-CoV-2 (National Institute for Biological Standards and Control, UK), was included as an assay control.
